# Integrative spatial transcriptomic analysis pinpoints the role of the ferroxidase, TaMCO3, in wheat root tip iron mobilization

**DOI:** 10.1101/2025.01.22.634289

**Authors:** Riya Joon, Gourav Singh, Deepshikha Tyagi, Varsha Meena, Vishnu Shukla, Kanupriya Agrwal, Shivani Saini, Mankiran, Hamida Banoo, Santosh Satbhai, Jagtar Singh, Terri Long, Eswarayya Ramireddy, Ajay K Pandey

## Abstract

Roots play an critical role in the sensing and absorption of essential minerals from the rhizosphere. Iron (Fe) deficiency, for example, triggers a well-known series of physiological and molecular responses within roots that facilitate uptake, which differs between monocots and dicots. In monocots, little is known about molecular responses that occur within specific root development zones in response to iron deprivation, and how these differences results in overall nutrient uptake. Here, we conducted a transcriptome analysis of wheat root tips under Fe deficiency (–Fe) and performed a comparative transcriptome analysis with the previous datasets generated from the whole root. Gene ontology analysis of differentially expressed genes highlighted the significance of oxidoreductase activity and metal/ion transport in the root tip, which are critical for Fe mobilisation. Interestingly, wheat, an allohexaploid species consisting of three different genomes (A, B, and D) displayed varying gene expression levels arising from the three genomes that contributed to similar molecular functions. Detailed analysis of oxidoreductase function at the root tip revealed multiple <u>m</u>ulti-<u>c</u>opper <u>o</u>xidase (MCO) proteins, such as Fe-responsive TaMCO3, that likely contribute to the overall ferroxidase activity. Detailed characterisation of *TaMCO3* shows that it complements the yeast FET3 mutant and rescues the –Fe sensitivity phenotype of *Arabidopsis atmco3* mutants by enhancing vascular Fe loading. Transgenic wheat lines overexpressing TaMCO3 exhibited increased root Fe accumulation and improved tolerance to –Fe by augmenting the expression of Fe-mobilizing genes. Our findings highlight the role of spatially resolved gene expression in –Fe responses, suggesting strategies to reprogram cells for improved nutrient stress tolerance.

## INTRODUCTION

Iron (Fe) is an essential micronutrient for plant growth development and plays a key role in regulating numerous cellular processes. Although Fe is present in the soil, it is primarily present in the insoluble Fe^3+^ form (Krohling et al., 2016), which is not readily available for plant uptake. Low soil Fe availability directly impacts plant productivity, with the onset of chlorosis, decreased root elongation, and reduced seed yield (Guerinot, 2007). Consequently, plants rely on multiple strategies to enhance Fe solubilization and uptake from soil. Recent research has provided new insights into Fe uptake *via* Strategy I (reduction) and Strategy II (chelation) mechanisms (Kobayashi and Nishizawa, 2012), suggesting that plants adapt by selecting modules from both strategies based on environmental needs (Li et al., 2023). For instance, dicots under alkaline soil conditions utilize a modified strategy I whereby Fe^3+^ reducing metabolites are secreted to facilitate Fe^2+^ uptake despite high pH and carbonate presence (Li et al., 2023).

Non-graminaceous plants, such as *Arabidopsis*, use a reduction-based strategy (Strategy I) in which plasma-membrane (PM)-localized H^+^-ATPases (AHAs) release protons to increase rhizosphere acidification and promote Fe^3+^ solubility. Subsequently, Fe^3+^ is reduced to the more soluble ferrous Fe^2+^ by ferric reduction oxidases (FROs) at the apoplast, and Fe^2+^ is transported into the root *via* the iron-regulated transporter (IRT1) (Santi and Schmidt, 2009). Graminaceous plants, including wheat, barley, and maize, utilize a chelation-based strategy (Strategy II). In this Strategy, plant roots secrete phytosiderophores (PS), forming Fe^3+^-PS complexes in the rhizosphere. Fe^3+^-PS is then taken up into root cells through the yellow stripe (YS) or yellow stripe-like (YSL) transporters for further long-distance transport (Curie et al., 2009; Kobayashi et al., 2010). The ability of plants to respond to soil nutrient availability is of fundamental importance for their adaptation to the environment. Changes in root architecture can mediate the plant adaptation to soils in which nutrient availability is limited by increasing the total absorptive surface of the root system. Under Fe-deficiency (–Fe) conditions, roots can also mobilize Fe by releasing specialised, complex metabolites such as coumarins, PS and their derivatives, which facilitate Fe mobilization (Yi and Guerinot, 1996; Fourcroy et al., 2014; Robe et al., 2021).

In general –Fe largely results in decreased root growth and reduced biomass thereby affecting the overall architecture (Wang et al., 2022). The molecular changes under –Fe in roots were investigated in different plant including Arabidopsis, tomato, rice, wheat and maize, (Li et al., 2016; Zanin et al., 2017; Sun et al., 2017; Martín-Barranco et al., 2020; Chen et al., 2022; Gabay et al., 2023; Molnár et al., 2023). Altogether, it was noted that the dynamic changes in the gene expression pattern remained temporal-specific yet specific transporters were identified that control Fe translocation between the subcellular compartments (Yokosho et al., 2009; Conte and Walker, 2011). The root tip is the first part of the root that comes into contact with the changing dynamics of soil and is expected to play a critical role under nutrient stress. Therefore, the root apex represents the most probable site for sensing Fe limitation and excess (Baluška et al., 1990). Accumulating evidence suggests that, compared to other regions of the root, the root tip is the most sensitive site for Fe fluctuation (Zhang et al., 2011; Zhang et al., 2012; Li et al., 2016). Although previous studies have described the impacts of –Fe on roots, many questions remain about root tip-specific responses in monocots.

Once Fe is taken up by the root epidermal cells it is radially transported to the vasculature *via* the symplastic pathway and exported into the xylem vessels in the form of Fe^2+^. It has been speculated that ferroportin (FPN)-like protein 1 is involved in this process (Morrissey et al., 2009). At this stage, it is oxidized from Fe^2+^ to Fe^3+^ by plasma membrane-localized ferroxidases. Arabidopsis root tips exhibit high ferroxidase activity, mediated by *Arabidopsis thaliana* Low Phosphate Root 1 (AtLPR1), a multicopper (MCO) ferroxidase, which facilitates Fe mobilization under phosphate (Pi) deprivation (Naumann et al., 2022; Xu et al., 2022), as well as callose deposition (Müller et al., 2015). Similarly, OsLPR5 plays a significant role in plant growth and development in rice by regulating a subset of molecular components involved in Pi-homeostasis that control lateral root growth and grain filling (Ai et al., 2020). At this stage Fe is transported as Fe^3+^-citrate or Fe^3+^-malate complexes. Recent studies in Arabidopsis have shown that the casparian strip selectively regulates nutrient absorption, blocking unregulated movement and enforcing symplastic or transcellular pathways. In the root tip, the polar localization of transporters aids in nutrient uptake by positioning specific transporters for efficient uptake (Robe and Barberon, 2023). Thus, specific plant tissue types facilitate the Fe uptake, transport, storage, and redistribution and maintain a balance to ensure optimal physiological functions and prevent toxicity.

While there are increasing insights into the function of specific tissues and cell types in model species, little is known about how root tip responses in crop species. Hexaploid wheat is an important source of energy and nutrition in many countries. Previously, the genome-wide transcriptome generated from whole root under –Fe (after 4, 8 and 20 days) revealed differential expression of numerous genes, including those encoding membrane transporters, metabolic pathway genes and transcription factors those are crucial for maintaining Fe homeostasis (Kaur et al., 2019; Kaur et al., 2023). In the current study, we specifically examined wheat transcriptional responses in the root tip after 8 days of –Fe. Using a comparative spatial transcriptomics approach, we identified the wheat ferroxidase candidate gene, TaMCO3, and characterized its role in Fe mobilisation and deficiency response. Altogether, this work highlights the importance of tissue-type specific –Fe responsive genes in the root tips of hexaploid wheat that could serve as targets for modifying cellular behaviour under nutrient stress.

## RESULTS

### Root tip transcriptome analysis identifies “representative-core –Fe response”

Root tips (until to the meristematic zone) are a primary site of Fe-sensing (Zhang et al., 2018). Therefore, we performed RNAseq analysis of wheat root tips (crown, seminal and primary roots) exposed to optimal +Fe (control) and –Fe (post 8 days) to gain further insight into how these conditions impact transcriptional responses. Perls/DAB staining **(Figure 1A)** validated the low accumulation of Fe in the root tips of –Fe treated plants, compared to control treatments. We were able to map more than >90% of the reads generated in the transcriptome. We observed a wide dispersion of gene expression from the two treatments **(Table S1),** with the highest Log_2_Fold-Change (Log_2_FC) ranging from +8.0 (upregulated) to -6.6 (downregulated). A total of 6234 differentially expressed genes (DEGs) were identified in response to –Fe (**Table S1)**. Out of this 4805 genes were upregulated, and 1429 genes were downregulated (LogFC >1.0) **(Figure 1B)**. Volcano plot suggests statistical significance (P value) distribution versus magnitude of change as LogFC **(Figure 1C)**. Gene Ontology (GO) enrichment analysis Differentially Expressed Genes (DEGs) revealed enrichment of biological processes related to metal ion transport and other cellular responses **(Figure 1D, Table S2)**.

**Figure 1:**
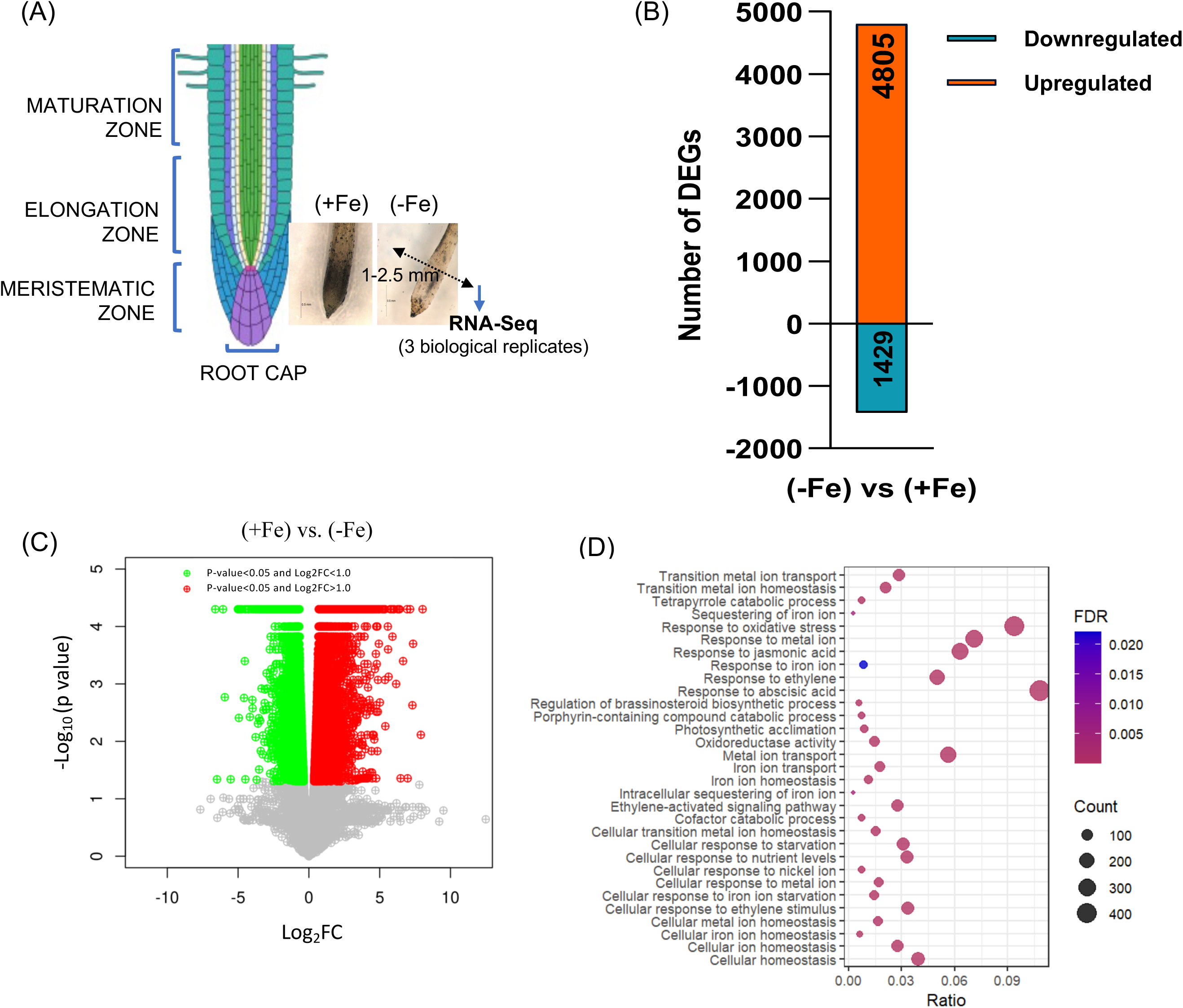
RNAseq analysis of the wheat root tip. **(A)** Schematic representation of the wheat root tip highlights the three distinct root zones: maturation, elongation, and meristematic zones. Lower right, microscopy images of root tips under both +Fe (iron-sufficient) and –Fe conditions (Perls/DAB staining), illustrating morphological differences in these zones. The region from which tissue was collected for subsequent RNA-seq analysis is marked on the diagram, corresponding to first 1-2.5mm of the root, which mostly includes the meristematic zone of the total root. Images were generated using BioRender. (B) Bar graph illustrating the total number of DEGs identified in root tip samples under Fe-deficient (-Fe) versus Fe-sufficient (+Fe) conditions through RNA-seq analysis. A total of 4,805 genes were upregulated, while 1,429 genes were downregulated in response to –Fe. **(C)** Volcano plot illustrating the differential gene expression between +Fe and –Fe root tip samples subjected to RNA-seq analysis. The plot highlights the statistical significance (y-axis, -log10 p-value) versus the magnitude of change (x-axis, log2 fold change) for each gene. Genes with significant upregulation under –Fe conditions are indicated on the right (red colour), while those downregulated are on the left (green colour). **(D)** Gene Ontology (GO) enrichment analysis for –Fe DEGs for “root tip-specific” responses. A total of 5036 DEGs were analysed.

### Comparative transcriptome analysis of root tip and whole root under –Fe

We then compared root tip transcriptional responses to that of our previously published transcriptomic dataset from whole roots exposed to 4 and 8 D (days) of –Fe (Kaur et al., 2023). Interestingly, we observed that 377 genes that are commonly expressed at all the data points and in root tips (**Table S1**); whereas 5036 DEGs were “root-tip specific” responses. Out of the total DEGs, 567 genes was found to be common between 8D root and root tip DEGs subjected to –Fe **(Figure 2A)**. In general, in both the data sets we found that genes encoding Nicotinamine acid synthase (NAS) and Nicotianamine aminotransferase (NAAT) proteins are highly induced by –Fe (**Table S2**). Various transporters, including Zinc-induced facilitator-like (ZIFL) protein-encoding genes known to be involved in PS release, were also highly upregulated. Additionally, YSL genes were highly represented in the RNAseq data set **(Figure 2B)**. These transcriptional changes in wheat roots were similar to the previous, wherein genes encoding for Fe acquisition and uptake were reported (Kaur et al., 2019; Kaur et. al., 2023). Multiple transcription factors, including putative orthologs of the basic-helix-loop helix (bHLH) family genes with a known roles in –Fe response, were also highly represented in our dataset **(Figures S1A** and **B)**. Overall, our data reveal that the wheat root tip transcriptional response includes well-established “core –Fe response” (Schmidt and Buckhout, 2011).

**Figure 2:**
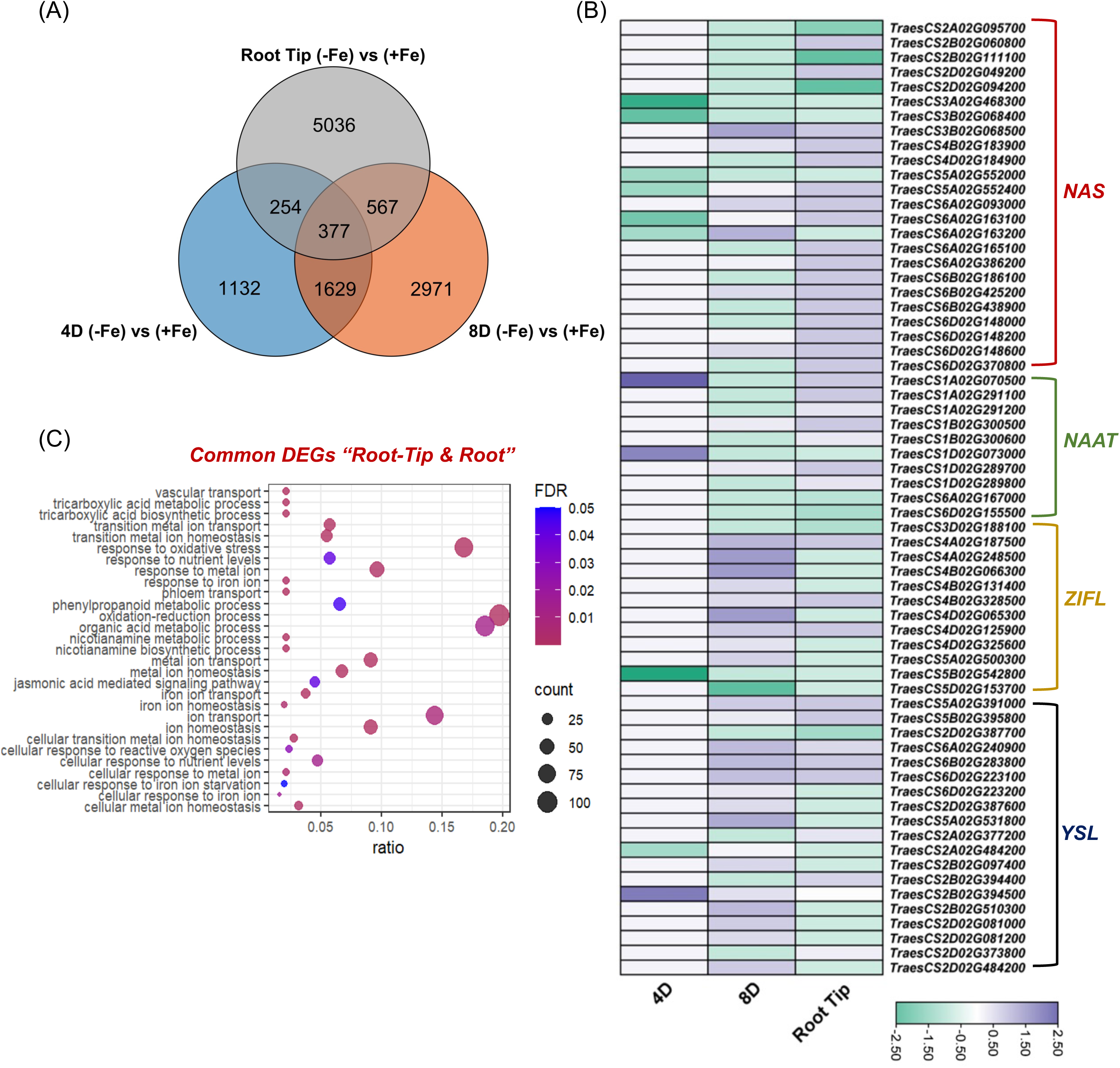
Comparative GO annotation of -Fe DEGs. **(A)** Venn diagram showing the overlap of upregulated genes among three conditions: root tip –Fe treated samples, whole root 4-day –Fe treated samples, and whole root 8-day –Fe treated samples. **(B)** Heatmap depicting the expression patterns of DEGs under –Fe (4 and 8 days) and root tip –Fe conditions. The colour scale represents the relative expression levels across the different conditions, illustrating both the upregulation and downregulation of these genes in response to –Fe. **(C)** Gene Ontology (GO) enrichment analysis for 567 common DEGs identified in both root tip (–Fe 8D) and whole root samples (–Fe 8D) under –Fe conditions.

For the root-tip, our analysis reveals enrichment of GO terms such as “response to oxidative stress”, “response to metal ions”, “metal ion transport”, and other related functions (**Figure 1D** and **Table S2**). Next, we performed a comparative analysis of GO annotation for the DEGs representative from root tip and whole root. A comparative GO annotation was also performed for the genes commonly identified between root-tip and root DEGs datasets (**Table S3**). The comparative GO annotation showed enrichment (ratio and number of DEGs) for terms such as “response to oxidative stress”, “ion transport”, “Fe transport”, “response to metals” and “organic acid metabolism” (**Figure 2C**). Although many of the GO terms overlapped among these datasets, in general, we observed a higher proportion of genes in the root-tip specific response when compared to the shared genes dataset for a similar biological process.

### Oxidoreductase activity is the major contributor at the root tip

Next, we checked the distribution of GO terms within the A, B and D genome transcripts. In general, we observed that although the number of genes varies among the homoeologs, the nature of GO functions was very similar between genomes (**Figure 3A and Table S4**). We observed that DEGs associated with the GO category, response to hormone, represented the largest contribution of DEGs across the sub-genome. It was notable that genes within the GO category “oxidation-reduction process” were highly represented by the transcript largely encoded by A and B genomes. This suggest that root tip are important site for redox related process to mobilize Fe in uptake. Further detailed analysis of this category of genes revealed many genes encoding for Laccases-like <u>M</u>ulti-<u>C</u>opper <u>O</u>xidases (MCOs). MCOs catalyse the oxidation of Fe, and are known to be involved in modulating root-to-shoot Fe partitioning (Bernal and Krämer, 2021). Heat map analysis of wheat MCOs suggested that a specific cluster of MCOs genes are differentially expressed in the root tip in response to 4 and 8 days of –Fe (**Figure 3B and Table S5**). In addition to this we observed differential expression of peroxidases encoding transcript but no expression was observed for ferric chelate reductase (**Figure S2A&B**). This analysis suggests that wheat MCOs are possibly involved in Fe mobilization in a cell-type specific manner *via* its ferroxidase activity.

**Figure 3:**
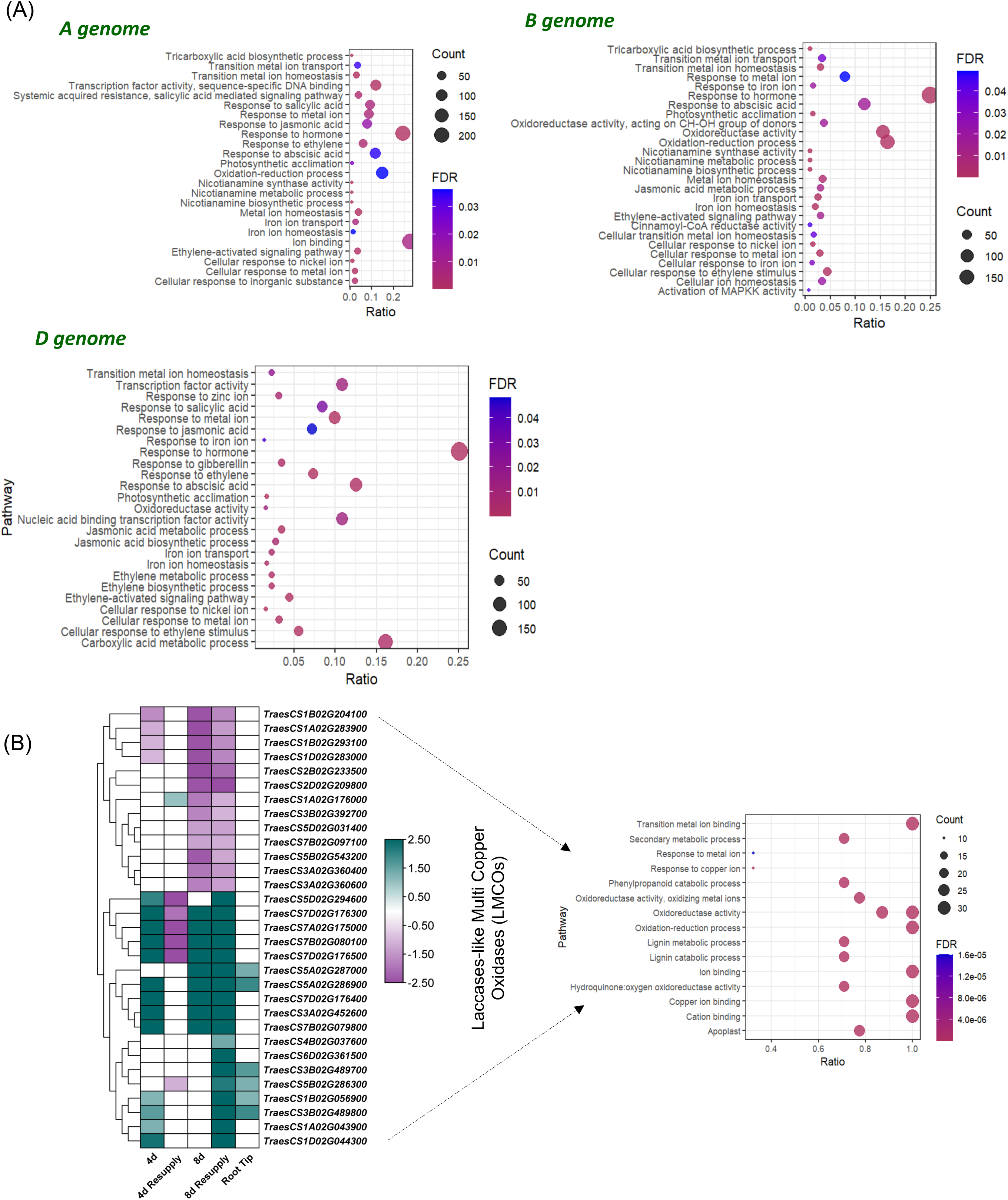
GO enrichment of the sub-genome transcripts and identification of –Fe induced wheat MCOs. **(A)** GO enrichment analysis was conducted for wheat homeologue-specific gene expression, revealing various pathways involved in the A, B, and D genomes of hexaploid wheat. **(B)** Left, heat map of the multi-copper oxidase (MCO) family, constructed based on RNA-seq data from wheat roots at 4 days and 8 days of –Fe (Kaur et al., 2023). Right, specific GO enrichment analysis focused on oxidoreductase activity in wheat root tips under iron-deficient conditions. The analysis illustrates the significance of oxidoreductase activity in the regulation of iron in wheat. The data highlight the upregulation of genes associated with oxidoreductase activity, underscoring their crucial role in iron homeostasis and adaptation to –Fe.

### *TaMCO3* expression analysis and Yeast *FET3* mutant complementation

In yeast, the multicopper oxidase FET3 operates as a ferroxidase, converting Fe²□ to Fe³□, which is essential for Fe absorption (Hoopes and Dean, 2004). To identify the closest ortholog of FET3 in wheat, we conducted an extensive phylogenetic analysis of genes induced by –Fe that encode MCOs **(Figure 4A)**. The percentage identify of FET3 with wheat MCOs ranges from 18-30% (**Figure S3**). The members of TaMCO of Clade I show the closest identity with the yeast FET3 protein. We also checked the expression of representative MCOs (*TraesCS6D02G361500*, *TraesCS7D02G176300, TraesCS1D02G044300*, *TraesCS3A02G452600, TraesCS5A02G286900* and *TraesCS4D02G035000)* from all the six clades. Our expression analysis under –Fe indicated that all the TaMCOs showed higher expression on 8D compared to 4D. Specifically, upon resupply of Fe, *TraesCS6D02G361500*, *TraesCS1D02G044300*, *TraesCS3A02G452600* and *TraesCS4D02G035000* show higher expression when compared to control or –Fe conditions (**Figure 4B**). We noted that two TaMCOs viz. *TraesCS7D02G176300* and *TraesCS5A02G286900* show significantly high expression at 4D of –Fe and remained unaffected by resupply at 8D. Among these candidates, TraesCS5A02G286900 was significantly upregulated in the root tip transcriptomic datatset (**Figure 3B**). Interestingly, *TraesCS5A02G286900* and *TraesCS4D02G035000* are in the same clade, along with ScFET3. Additionally, under the -Fe condition, *TraesCS5A02G286900* showed higher expression in the root tip than *TraesCS4D02G035000* **(Figure S4A)**. At the protein level, *TraesCS5A02G286900* shares 28.97% identity with ScFET3 and 49.56% with Arabidopsis At5G21105 (AtMCO3). While all the proteins include the typical conserved cupredoxin and ascorbate oxidase domains, both AtMCO3 and *TraesCS5A02G286900* lack the plasma membrane domain **(Figure S3 and Figure 5A**). Thus, we further evaluated *TraesCS5A02G286900* as a candidate ferroxidase encoding protein and referred to as *TaMCO3*.

**Figure 4:**
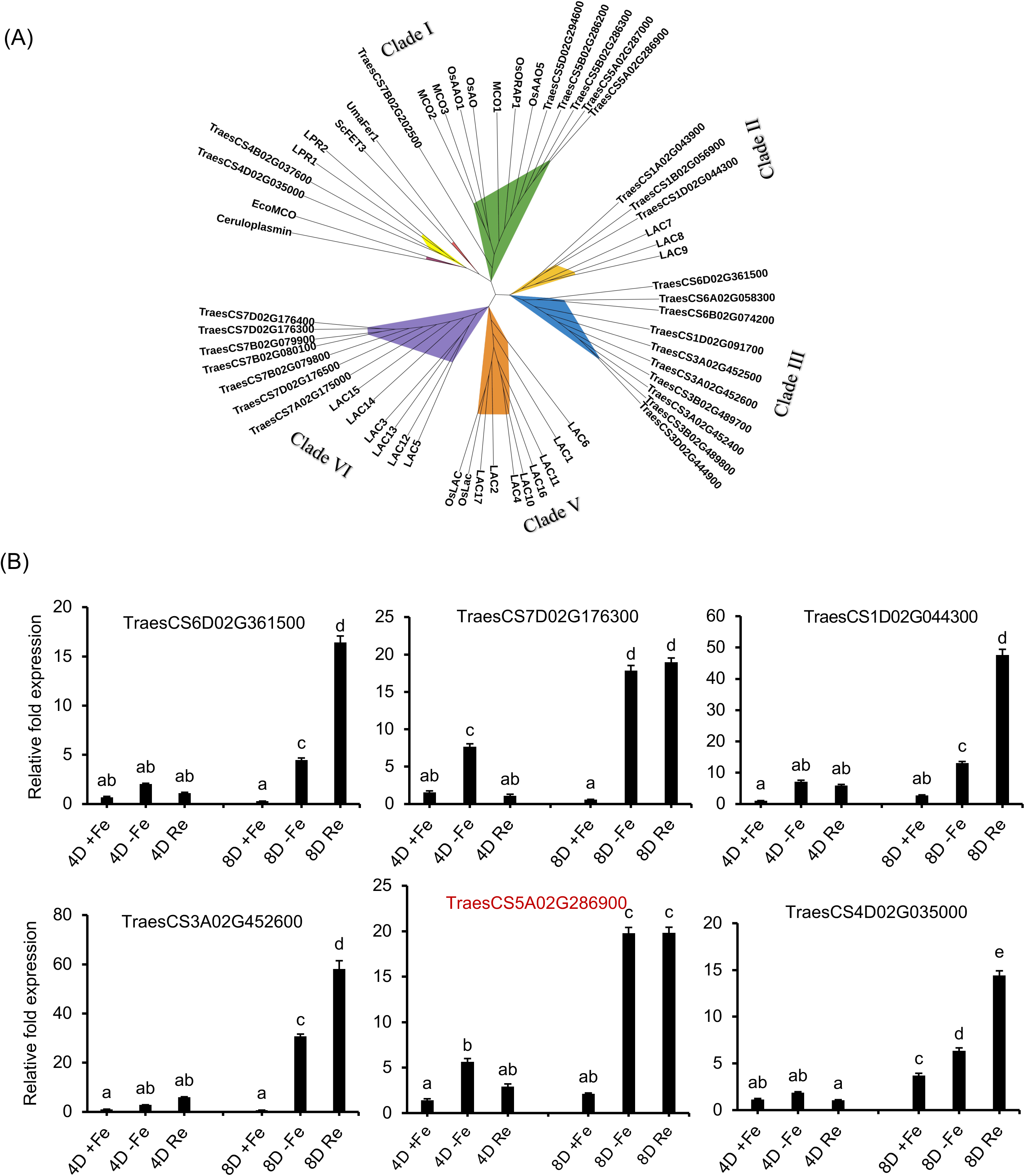
Expression and phylogenetic analysis of differentially expressed wheat MCOs. **(A)** Phylogenetic tree of Fet3-like genes, including ScFET3 (NP_013774), Uma Fer1 (XP_011386068), EcoMCO (EDU66368), Ceruloplasmin (EAW78883), AtMCOs (LAC1-17, LPR1-2, MCO1-3), and TaMCOs. MCO proteins were aligned using the MUSCLE algorithm, and an unrooted phylogenetic tree was generated using the neighbor-joining method with MEGA 11, incorporating a bootstrap value of 1000. **(B)** qRT-PCR analysis of *TaMCO* genes. Wheat seedlings subjected to different concentration of Fe-EDTA as +Fe (80 µM) and –Fe (1 µM) and resupply (Re) (80 µM) for 4 and 8 days under hydroponic conditions. Each bar indicates the mean of three replicates with the indicated standard deviation of the mean. C_t_ values were normalized using wheat ARF1 as an internal control.

**Figure 5:**
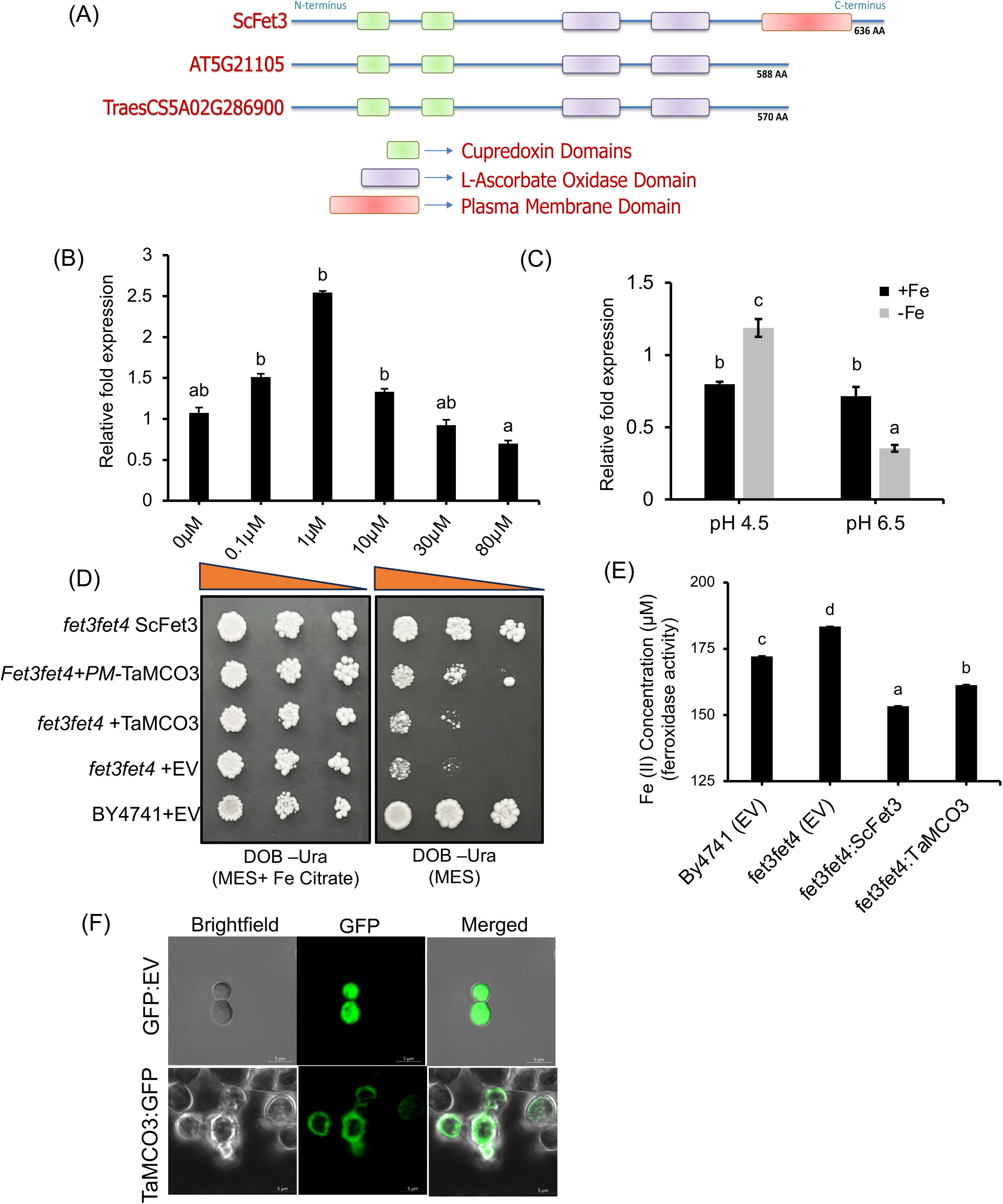
Characterization of wheat candidate ferroxidases**. (A)** Schematic of of ScFET3, AT5G21105 (AtMCO3) and TraesCS5A02G286900 (TaMCO3) domains **(B)** Expression of *TaMCO3* under various Fe dosages. **(C)** Expression of *TaMCO3* under different pH conditions, The expression was normalized using *TaARF1* as internal control, and fold change was calculated using the 2^−ΔΔCT^ method. Vertical bars indicate the standard deviation (n=3), and letters above the bars indicate significant differences (P<0.05) between the various tissues by one way ANOVA test. **(D)** A yeast complementation assay was performed with the yeast *fet3fet4*Δ mutant, which exhibited growth impairment compared to the wild-type BY4741. However, growth was partially restored upon the addition of TaMCO-PM. Fe(II) disappearance in yeast total protein was measured using ferrozine as the chelator. **(E)** Ferroxidase activity indicated by plotting Fe(II) disappearance, with total protein isolated from yeast cells. **(F)** Wheat MCOs were cloned into the pRS315TEF_GFP vector for localization studies in *Saccharomyces cerevisiae*. Images were acquired using LEICA DM600CS confocal microscope and processed with Fiji-ImageJ. GFP fluorescence was detected at 500– 530 nm, for both pRS315TEF_GFP empty vector and pRS315TEF_GFP:TaMCO3 with magnification 25x and 40x.

Next, we examined the expression of *TaMCO3* in roots under conditions of varying Fe availability and found increasing fold change in transcript levels under deficient conditions (0, 0.1, and 1 µM Fe) when compared to the control (80 µM Fe). This was inconsistent with our previous qRT-PCR analysis (**Figure 4B**). We also observed that the expression of *TaMCO3* decreased with increasing Fe levels, suggesting its expression response is as a result of a change in Fe in roots. This highlights that the expression response to *TaMCO3* depends on the Fe availability **(Figure 5B)**. Furthermore, considering the possible redox fluctuations in roots driven by pH (Husson, 2013; Zhang et al., 2015), we assessed *TaMCO3* expression levels at various pH conditions. The gene showed greater expression at low pH 4.5 compared to a more neutral pH of 6.5, likely due to redox shifts that influence Fe availability in the rhizosphere **(Figure 5C)**. We also tested the effect of protein synthesis inhibitor cycloheximide (CHX) on the expression of *TaMCO3*. In contrast, the expression of TaMCO3 was high in the presence of CHX in both +Fe and –Fe conditions. This suggests that the presence of CHX prevents the low expression of *TaMCO3* (Figure S4B), supporting the idea that *TaMCO3* expression relies on newly synthesised transcription factors (de novo TFs).n ExVip expression database (Ramírez-González et al., 2018) at different Zadosky scales (Z), indicated high expression of *TaMCO3* in the roots (Z13 and Z39 stage) compared to other wheat tissues (**Figure S4C)**. A previous study has analysed wheat root tip transcriptome in monocot-specific *12-OXOPHYTODIENOATE REDUCTASE* (OPRIII) overexpression lines that were shown to modulate key differences in wheat root architecture (Gabay et al., 2023b). The expression *TaMCO3* was high in the transgenic lines (**Figure S4D**). The expression of *TaMCO3* was also checked during grain development and grain tissue. Our qRT-PCR expression data indicated the highest expression at 28 days after anthesis (DAA), with the highest in pericarp tissue (**Figure S4E**).

For functional characterization, we investigated if TaMCO3 (A homoeologue: TaMCO3-5A) could compensate for the functionality of yeast FET3. We fused the open reading frame (ORF) of TaMCO3 to the plasma membrane (PM) anchoring domain of yeast FET3. In yeast, the PM-anchored TaMCO3 successfully alleviated the Fe sensitivity phenotype of the fet3 mutant (Brun et al., 2022), localizing to the plasma membrane and restoring ferroxidase activity. Our complementation suggested complementation of FET3 activity in yeast by TaMCO3-5A (**Figure 5D**) was able to complement the ferroxidase activity in the yeast (Figure 5E). The TaMCO3-5A:GFP was found to be localised to the plasma membrane (**Figure 5E**). Thus, our data indicates that TaMCO3 serves a similar role in Fe oxidation as FET3 does in yeast and TaMCO3 with ferroxidases activity may be root-specific.

### TaMCO3 complements Arabidopsis mutant and contributes to –Fe tolerance

To investigate the subcellular localization of TaMCO3, we transiently expressed TaMCO3-YFP in *Nicotiana benthamiana* leaves using *agrobacterium*-infiltration. Our findings indicated that the TaMCO3 fusion protein primarily localized to the periplasmic space and the plasma membrane in tobacco leaf cells **(Figure 6A)**. This localization was supported by similar results obtained in yeast cells, where TaMCO3 also localized to the plasma membrane **(Figure 5E)**, suggesting that subcellular targeting may be conserved across species. In addition to the localization study, we assessed the functional activity of TaMCO3 in relation to *Fe deficiency*.

**Figure 6:**
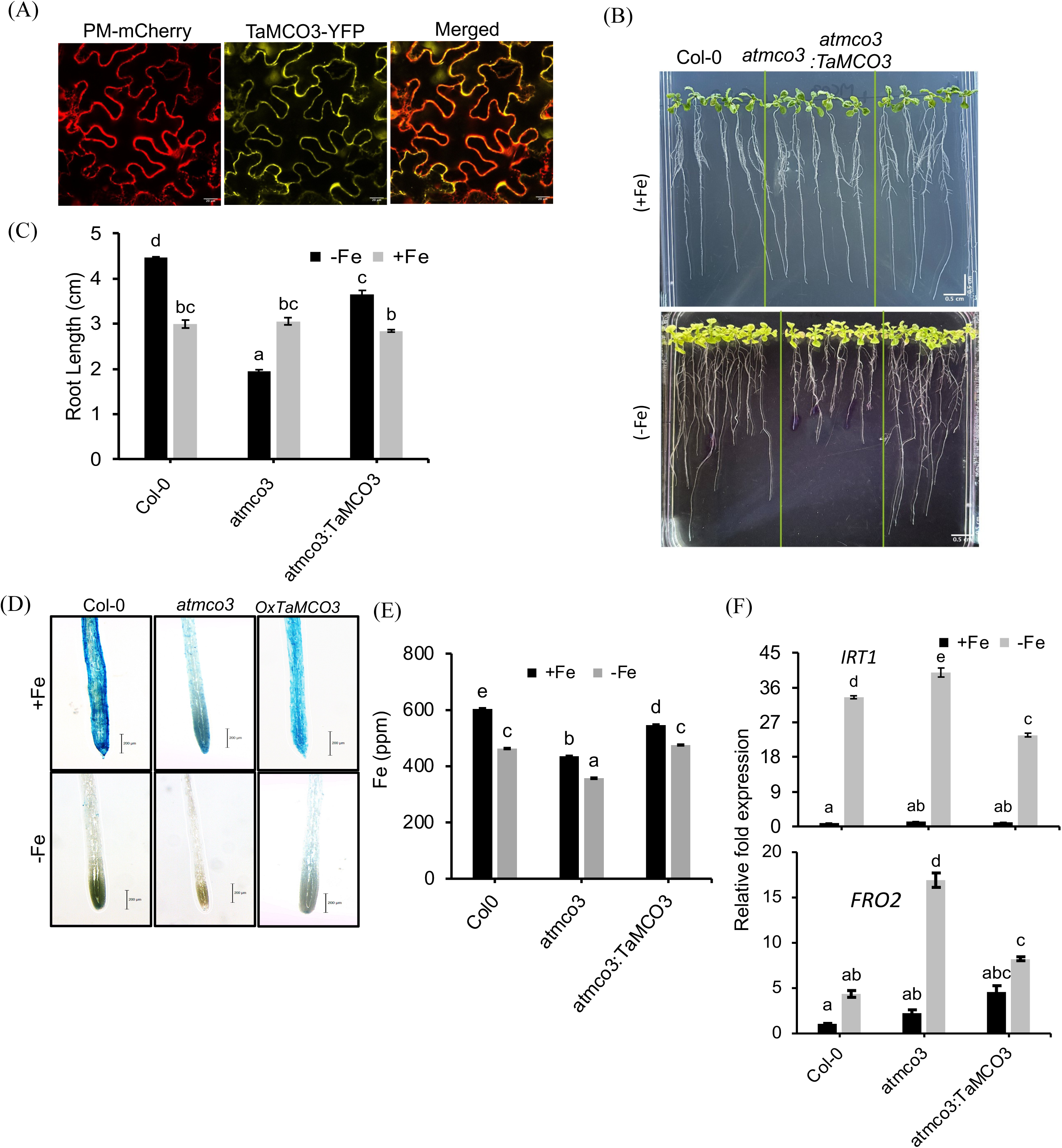
Localization of TaMCO3 and complementation of *atmco3* with TaMCO3**. (A)** Wheat MCOs localization in agro-infiltrated *Nicotiana benthamiana* leaves. Images were acquired 48 hours post-agroinfiltration using a LEICA DM600CS confocal microscope and processed with Fiji-ImageJ. Red fluorescence was detected at 580–630 nm. Left, plasma membrane marker pMRK-RFP, middleTaMCO3-YFP, right, overlay at magnifications of 25x. **(B)** Overexpression of TaMCO3 in an *atmco3* mutant background, demonstrating rescue of the Arabidopsis knockout mutant *atmco3* phenotype. Plants were grown on Hoagland solid medium for 5 days, then transferred to +Fe (300 µM FeEDTA) and –Fe (300 µM Ferrozine) media for 2 weeks. **(C)** Root lengths measured using ImageJ software (Version 1.38). The values presented are representative of seven independent biological replicates. Letters above the bars indicate significant differences (P < 0.05) between genotypes, as determined by a two-way ANOVA test. **(D)** Representative images of iron distribution in Col-0, *atmco3*, and TaMCO3 overexpressing Arabidopsis lines, detected by Perls staining, Seedlings were germinated on Hoagland agar media for 5 days and then transferred to Hoagland minimal media amended with +Fe (80 µM FeEDTA), −Fe (300 µM Ferrozine), for 10 days. **(E)** Mean iron content, quantified by ICP-MS, error bars representing standard deviation. Letters above the bars indicate statistically significant differences (P < 0.05) between genotypes, as determined by one-way ANOVA. **(F)** Expression analysis of *AtIRT* and *AtFRO2* in Col-0, *atmco3* and *TaMCO3* overexpressing lines.

Mutations in multicopper oxidases (MCOs) in *Arabidopsis* increase sensitivity to –Fe (Bernal and Krämer, 2021). Thus, we proposed that TaMCO3 could play a pivotal role in Fe mobilization and uptake in conditions of Fe scarcity. To evaluate this hypothesis, we generated multiple *atmco3* lines overexpressing MYC:TaMCO3. Two independent transgenics were confirmed for fusion protein expression (MYC:TaMCO3) by protein blotting and were used for further characterization **(Figure S5)**. Our phenotypic assessment showed that under +Fe, the root lengths of Col-0, *atmco3* mutants, and the TaMCO3-complemented lines exhibited no significant differences **(Figure 6B and Figure S5)**. However, under –Fe, the *atmco3* mutants displayed a significant decrease in root growth, indicating an increased sensitivity to low Fe levels. Across different TaMCO3 complemented lines, the root lengths were comparable to those of Col-0, suggesting that the overexpression of TaMCO3 effectively rescued the –Fe phenotype in the *atmco3* mutant **(Figure 6B and C, Figure S5).**

To further examine the role of TaMCO3 in Fe accumulation, we quantified the Fe content in the roots of these Arabidopsis lines using inductively coupled plasma mass spectrometry (ICP-MS). Our results demonstrated that the TaMCO3 overexpressing lines were capable of restoring Fe accumulation in the roots to levels similar to those of Col-0, whereas the *atmco3* mutants exhibited significantly lower Fe accumulation **(Figure 6D and E)**. This indicates that TaMCO3 is engaged in the Fe uptake process in wheat, and its overexpression in Arabidopsis can compensate for the functional loss in the *atmco3* mutant.

Additionally, our results were supported by Perl’s staining, which specifically detects Fe³□ in plant tissues. The staining patterns revealed a significant restoration of Fe uptake in the TaMCO3 complemented lines, while the *atmco3* mutants exhibited reduced Fe uptake, further reinforcing the role of *TaMCO3* in Fe homeostasis **(Figure 6D)**. Under –Fe condition the expression of wheat *IRT1* and *FRO2* in TaMCO3 complemented lines showed similar expression response as observed in Col-0 **(Figure 6F)**. Overall, these findings suggest that TaMCO3 plays a crucial role in Fe mobilization and uptake, particularly under –Fe conditions, thereby helping to alleviate symptoms of –Fe in both Arabidopsis and wheat.

### Overexpression of TaMCO3 contribute tolerance to –Fe

Next, we assessed wheat root ferroxidase activity by growing plants in the presence of FeSO_4_ (as the only Fe source), and quantifying ferrous Fe chelation to ferrozine as a proxy for ferroxidase activity. Our results show that within days of exposure to FeSO_4_ there is a decrease in ferrozine-Fe^2^ ^+^ complexes, (**Figure 7A**), suggesting high oxidation activity. We then tested the total ferroxidase activity in wheat roots in under Fe-sufficient and deficient conditions, and observed a significant disappearance of ferrozine-Fe^2^ ^+^ complexes in roots subjected to –Fe (**Figure 7B**). Next, to assess its function, we overexpressed TaMCO3 in wheat *cv*. Fielder using a constitutive promoter. Immature embryos were used to generate wheat transgenic lines (**Figure S6A**). We generated multiple transgenic lines, and characterized the two independent transgenic lines with the highest expression (**Figure S6B:** Line#2, #4). Immunoassay and localization studies of the transgenic lines expressing c-MYC:TaMCO3 suggested its presence at the root plasma membrane and was consistent across the independent transgenic lines This localization (**Figure 7C &Figure S7**).

**Figure 7:**
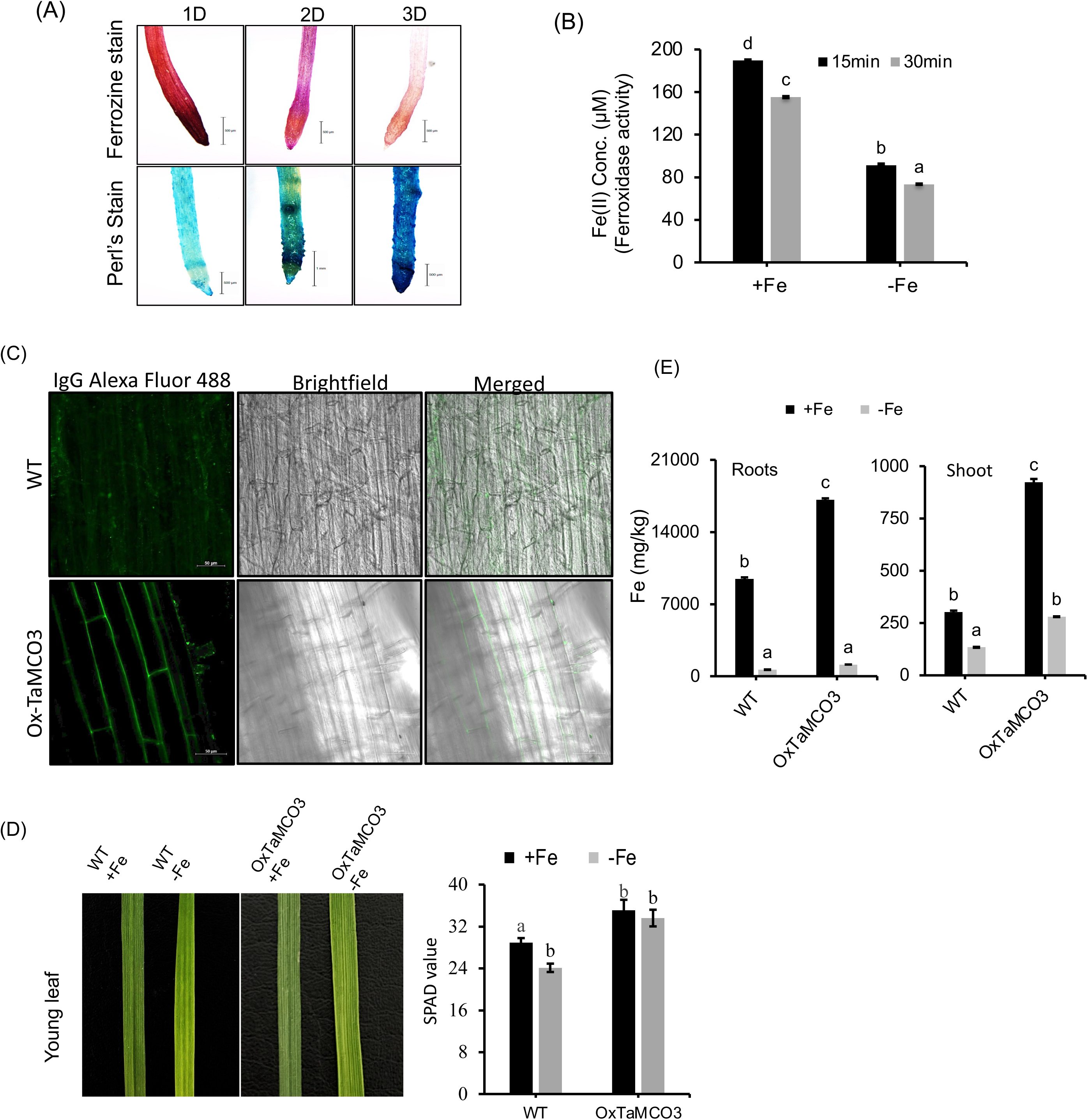
TaMCO3 expression in WT and the impacts of TaMCO3 overexpression in wheat. **(A)** Ferrozine and Perls staining of wheat root tips was performed after their growth in Fe□SO□-supplemented media for 1, 2 and 3 days (D). **(B)** Fe^2+^ concentration (in μM) measured after treating FeSO□ substrate with total root protein under control (80 μM Fe-EDTA) and –Fe (1 μM Fe-EDTA) conditions. The assay included three biological replicates and three technical replicates. The Fe^2+^-ferrozine complex formation was measured at 560 nm absorbance. Decreased absorbance of the Fe^2+^-ferrozine complex in the iron-starved samples indicates increased oxidation of Fe^2+^ in the media. **(C)** Whole-mount immunolocalization of leaf tissue. Antibody-treated (anti-c-Myc tag Mouse Monoclonal antibody) sections were transferred to microscopic slides, and images were captured in replicates at 10X using an Zeiss LSM 880 Confocal Laser Scanning Microscope. **(D)** Leaf SPAD (Konica Minolta SPAD-502, Japan) values of WT and TaMCO3 overexpression line under +Fe and –Fe conditions, seedlings (n = 28-30) were used for each treatment and in each seedling the fully expanded youngest leaves were used, with 3-4 spots per leaf. **(E)** The total iron content of the root and shoot was measured using ICP-MS.

Next, we examined the transgenic lines for response to –Fe. We found that TaMCO3 overexpression causes delayed young leaf chlorosis compared to WT, as evidenced by a decrease in SPAD values **(Figure 7D)**. Interestingly, overexpressing lines show no change in root architecture whereas the WT shows reduced root growth under –Fe conditions **(Figure S8A)**. Under control conditions, higher Fe levels were also observed in transgenic lines in both the root and shoot when compared to the non-transgenic (WT) **(Figure 7E and Figure S8C)**. Surprisingly, under –Fe conditions, both transgenic and WT lines show no significant differences of Fe; whereas, in shoots, significantly high Fe levels were measured in the transgenic lines. We speculate that elevated Fe in the shoots of these overexpression lines is due to increased translocation from the root to the shoot due to high ferroxidase activity. We also checked the expression of known Fe responsive genes. Our analysis suggests that while none of our candidate genes are differentially expressed under +Fe, during –Fe treatment, *TaYS1A* expression was elevated in *TaMCO3* overexpression transgenic lines compared to WT. No significant changes in the expression of *TaIRT1*, *TaIRO3*, *TaFIT, and TaNAS3* was observed in the transgenic lines subjected to –Fe condition. Interestingly, we observed high expression of *TaZIFL4.1*, *TaIRT1*, *TaIRO3*, *TaFIT,* and *TaNAS3* in WT under –Fe when compared to +Fe **(Figure 8A)**. This suggests that high uptake of Fe and oxidation in the root pericycle contributes to delayed chlorosis in TaMCO3-transgenic wheat.

**Figure 8:**
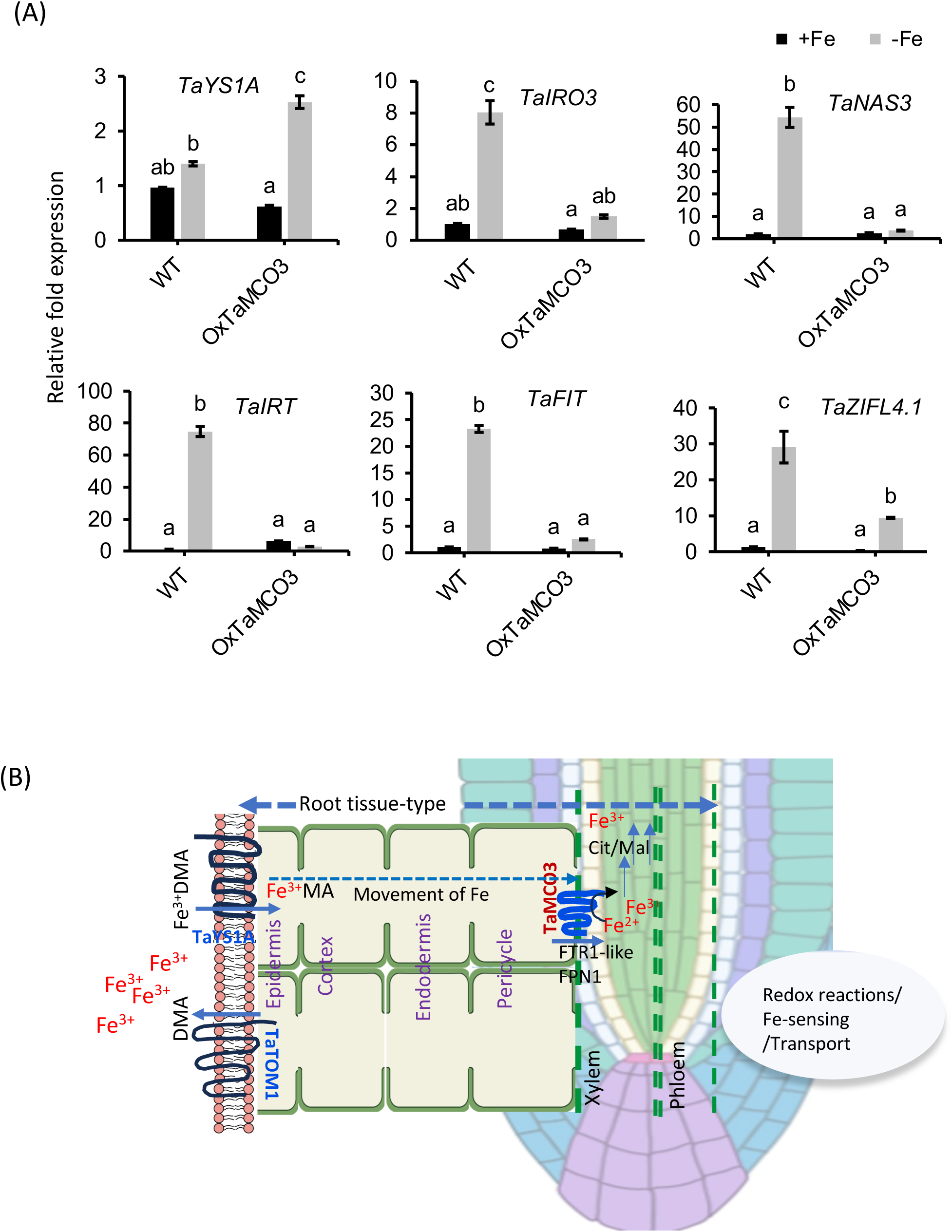
Expression analysis of known −Fe responsive genes and model summary. **(A)** Expression analysis of known Fe homeostasis genes in *TaMCO3* overexpressing lines. **(B)** Model summarising the key molecular and biochemical mechanisms uncovered in this study, highlighting the role of TaMCO3, a wheat multicopper oxidase, in mediating iron uptake and mobilisation. Known TOM (transporter of mugineic acids; Nozoye et al., 2010;) proteins assist in DMA (2′-deoxymugineic acid; Zhang et al., 2022) efflux of a major PS in wheat. Once the Fe-Chelates (Fe-PS) complex are formed they are subsequently taken up into the epidermis by TaYS1A and transported to the pericycle region. Fe^2+^ is then oxidised by ferroxidases (MCO such as TaMCO3) to Fe^3+^, which is complexed with citrate/malate, and transported via the xylem to the shoot. Images were generated using BioRender.

## Discussion

The root tip is a sensory hub that detects environmental signals, including nutrient fluctuations. It enables plants to adapt to varying soil conditions by altering root growth direction or rate (Jiang et al., 2016; Zhang et al., 2018) via transcriptional and translational regulatory processes that control root tropisms, cell polarity, and cell fate (Baluška et al., 1990). previous transcriptomic analysis of wheat roots subjected to –Fe identified genes involved in Fe mobilization and uptake (Kaur et al., 2019; Kaur et al., 2023). Organic acids such as citrate and malate were also shown to accumulate in high amounts in roots exposed to Fe deficient conditions (Kaur et al., 2019). In addition, the amount of PS released by the root tips plays a critical role in –Fe tolerance in hexaploid wheat (Meena et al., 2024). However, these studies have primarily focused on the total root response, thereby limiting our understanding of the key sensing site of the wheat root.

### Root tips are redox site for –Fe response

Root tips are likely the primary site for detecting and responding to heterogenous Fe fluctuations in the rhizosphere (Li et al., 2016). Here, we focus on understanding the molecular mechanisms of how hexaploid wheat root tips respond to –Fe. Our transcriptomics analysis of the root tip provides evidence that this region represents the site of core –Fe sensing and response **(Figure 2A)**. Our analysis also indicated root tip-specific GO annotations, some of which are important for Fe/metal mobilization and oxidoreductase activities **(Figure 1D)**. The presence of specific Fe forms in soils is primarily governed by soil conditions, including pH and redox status. Soil Fe is mainly present in the two most common oxidation states, Fe^2+^ and Fe^3+^, which interact with other elements and minerals to determine their availability to plants. Under nutrient deficiency conditions plant root tips maintain homeostatic redox balance to coordinate normal development. The overall redox status of specific roots zones under nutrient-limiting conditions or during abiotic stresses helps plants to cope with stresses. For example, in Arabidopsis, varying redox potentials have been observed in the root zones undergoing salinity stress (Jiang et al., 2016). In this study, we observed a large set of genes encoding for GO annotations related to the oxidoreductase/oxidation-reduction activity/process **(Figure 3B**) suggesting that Fe stress in root tip primarily relies on cellular redox activities.

Root cell-type specific transcriptional responses to –Fe have been reported previously (Dinneny et al., 2008). However, such cell-type-based studies are currently limited to the model plant Arabidopsis (Kajala et al., 2021). Recent advances in sequencing technologies has also enabled root tip single-cell transcriptomic analysis of hexaploid wheat under normal growing conditions (Zhang et al., 2023). Thus, we overlayed our root tip transcriptome data to this previously reported dataset, which was assembled using marker lines identify, class, and analyse distinct cell populations. Our analysis indicated that the spatial mapping of marker genes from our DEGs corresponds to 20 out of 23 previously reported cell types (Figure S9 and Table S6). This suggests a high confidence level in our dataset for uniformly representing these cell types. Subsequent analysis shows that 2273 transcripts differentially expressed in response to –Fe were mapped to different cell types, as reflected in the UMAP plot **(Figure S9B)**. Next, we checked the distribution of the mapped cluster DEGs among the cell-type. We observed that the most significant percentage (21.1 %) of genes were differentially expressed in the pericycle in response to –Fe, followed by the root hairs (11.6 %) and root border cells (10.6 %) (**Figure S10A and Table S7**). The pericycle is a transcriptional regulatory center for the –Fe response (Long et al., 2010; Gouran and Brady, 2024). Fe efflux transporters, including ferroportin (FPN1), are also localized on the plasma membrane of the pericycle, thus assisting the Fe^2+^ loading into the xylem (<u>Morrissey et al., 2009</u>). Multiple ferroxidases encoding proteins are also known to be localized in this vasculature tissue (Ai et al., 2020; Zhu et al., 2022). Although one must be cautious about comparing such datasets due to the difference in the root anatomic structures for monocots and dicots (Gouran and Brady, 2024), our analyses reinforces the importance of pericycle in Fe homeostasis.

### The spatial distribution of Fe DEGs reveals the role of ferroxidases

Our investigation revealed that redox homeostasis processes is one of the most prevalent GO categories associated with genes differentially regulated by –Fe in the wheat root tip (**Figure 3B**). This led us to identify multiple MCOs encoding ferroxidases potentially involved in efficient Fe mobilisation. In the wheat genome, natural variations in laccase genes have been shown to influence wheat–arbuscular mycorrhizal symbiosis (Zhong et al., 2023). Previous studies have also highlighted the role of TaLACs in wheat resistance to *Fusarium graminearum*, suggesting their potential to enhance stress tolerance (Sun et al., 2022). However, the current study identifies TaMCO3 as uniquely responsive to –Fe, with no detectable expression under other stress conditions. Recent advancements in single-nucleus RNA sequencing (snRNA-seq) and single-cell RNA sequencing (scRNA-seq) from wheat grain and root tissues—spanning over 20 distinct cell types—also did not reveal TaMCO3 expression (Zhang et al., 2019; Liu et al., 2021; Zhang et al., 2021; Shahan et al., 2022). This specificity to Fe stress makes TaMCO3 a particularly intriguing candidate for further investigation into its role in Fe homeostasis and plant.

Previously, plant laccase-like MCOs were characterized in detail for the first time in Arabidopsis. A total of 17 laccase-like multi-copper oxidases (MCOs) have been identified, each exhibiting diverse expression patterns. These patterns are either tissue-specific or associated with distinct stages of plant development (Turlapati et al., 2011). Their emerging role in nutrient homeostasis has been linked with enhanced ferroxidase activity. For example, OsLPR5, which exhibits ferroxidase activity, was shown to influence the growth and development process, including lateral root growth and grain filling, by influencing a subset of molecular components involved in Pi-homeostasis. This finding points to the distinct role of redox cycling activity mediated by MCOs to adapt –Fe stress. (Ai et al., 2020). Fe is transported as Fe^3+^-citrate or Fe^3+^-malate complexes. Thus, the ferroxidase activity of plant laccase-type MCOs, such as LPR5, helps maintain Fe and Pi homeostasis and adapt to nutrient stress by enabling Fe redox cycling (Hoopes and Dean, 2004; Zhu et al., 2022). Human MCOs have been identified and characterized to be ceruloplasmin proteins that possess ferroxidase activity regulating the Fe turnover by oxidizing ferrous from Fe^2+^ to ferric Fe^3+^ (Vashchenko and MacGillivray, 2013). Similarly, in lower eukaryotes like yeast, the MCO, FET3, was demonstrated to be a ceruloplasmin homolog showing ferroxidase activity (Stoj and Kosman, 2003). At the functional level, the cell-surface activity of FET3 is required for high-affinity transport of Fe (De Silva et al., 1995). Our phylogenetic analysis of the MCOs shows a specific cluster responding to –Fe induced expression **(Figure 4A)**. A detailed analysis reveals that wheat gene ID *TraesCS5A02G286900* (TaMCO3) is closest to *Saccharomyces cerevisiae* FET3. Ferroxidases are known to be expressed largely at the pericycle membrane and are reported to generate Fe^3+^ from thereby facilitating the mobilization of Fe^3+^-chelate forms (Bernal and Krämer, 2021; Liu et al., 2022; Zhu et al., 2022). In our study we found that TaMCO3-YFP is localized to the plasma membrane and overexpression of TaMCO3 likely leads to increased ferroxidase activity. Consequently, TaMCO3 reduces chlorotic response and thereby indirectly modulates the expression of Fe acquisition genes when exposed to –Fe **(Figure 7D and Figure 8A)**. This finding correlates with the observed high shoot Fe in the transgenic lines compared to WT, thereby providing resistance to –Fe in the rhizosphere. High ferroxidase activities in rice is also linked to enhanced grain Fe content that could be attributed to the high mobilization capacity of Fe to the grain tissue (Gu et al., 2024). Recently, rice OsLPR5 was also shown to be involved in modulating salt stress and Fe homeostasis-related genes (Ai et al., 2020; Zhao et al., 2023). This interconnection with salt and Fe stress was attributed mainly to redox changes in the plant roots. Together, these findings suggest that TaMCO3 in wheat root tips may be important for oxidizing Fe^2+^.

Consequently, overexpression of TaMCO3 results in robust root growth even under – Fe conditions, compared to WT **(Figure S8)**. Previously, another MCO, LPR1 was linked to Fe-dependent callose deposition in response to low phosphate conditions (Müller et al., 2015). MCOs has been linked to the cell wall maintenance and overexpression leads to high Fe^3+^ in both roots and shoots (Bernal and Krämer, 2021). In our overexpressing lines TaMCO3 protein was expressed constitutively and localized within the plasma membrane of different cell types, leading to high accumulation of Fe in shoots and roots **(Figure 7C)**. Such high accumulation might be beneficial for enhanced Fe loading in the xylem, and subsequently into seeds using tissue-specific endosperm promoters fused to Fe-transporters, or for imparting –Fe tolerance to growing wheat seedlings.

Based on our findings, we propose a model reflecting the importance of the different bottleneck checkpoints for Fe loading in specific cell/tissue type (**Figure 8B**). Overall, the genes contributing to the strategy II mode of Fe uptake are constitutively expressed and the expression of these genes is high under –Fe condition. In addition, although the contributions of sub-genome-derived gene expression vary, they all contribute to similar biological processes. As evident from the results, the wheat MCO3 localized in the plasma membrane plays an important role in generating Fe^3+^ in order to mobilize Fe to shoots. Although our study was limited by analysing the root tip rather than cell-specific transcriptomic data, it demonstrates the effectiveness of integrating stress-specific data with existing cell-specific information to uncover a new regulator in the –Fe response..

## EXPERIMENTAL PROCEDURES

### Plant material and growth conditions

The hexaploid wheat cv. C-306 and Fielder, provided by Punjab Agricultural University (Ludhiana, Punjab), were used in this study. These cultivars were grown in the experimental field at National Agri-food Biotechnology Institute, Mohali, Punjab, India. Wheat seeds were stratified overnight in the dark at 4°C, then germinated on Petri dishes lined with Whatman filter paper. Endosperms were removed from 4-day old seedlings, which were subsequently transferred to PhytaBox™ containers, where they were cultivated using Hoagland’s nutrient solution. For –Fe, 1 µM Fe(III)-EDTA was used, while control plants (+Fe) received 80 µM Fe(III)-EDTA without altering other nutrient concentrations. The plants were grown in a controlled growth chamber set at 21±1°C, with 50–65% relative humidity, and a photon flux density of 300 µmol m^−2^ s^−1^, under a 16-hour light/8-hour dark photoperiod. Roots and shoots were harvested from three biological replicates after 8 days post-deficiency and immediately frozen in liquid nitrogen for subsequent analysis. For RNAseq analysis, root tips (containing the meristematic zone) were collected up to ∼2.5mm from the apex. Total RNA was extracted from the tissue from three biological replicates and processed for transcriptome analysis as described earlier (Kaur et al., 2019; Kaur et al., 2023). For cycloheximide (CHX) treatment (Meiser et al., 2011), plants grown hydroponically were pre-treated with CHX 50 µM for 1 h to inhibit new protein translation. For Fe treatments, Fe starvation (−Fe) was induced using 1 µM Fe(III)-EDTA, while control (+Fe) and excess Fe (++Fe) conditions were maintained with 80 µM and 200 µM Fe(III)-EDTA, respectively, for 6 h. For dosage experiments roots were subjected to different concentrations of Fe and root were collected post 4 days of −Fe. A total of three experimental replicates (each containing 10–12 seedlings) were used for total RNA extraction from whole root or root tips.

### Bioinformatic and GO enrichment analysis

Transcriptome analysis was conducted as described earlier (Kaur et al., 2019; Kaur et al., 2023). Data was generated from three biological replicates for each treatment. Trimmomatic 0.35 was used to trim and filter reads, discarding those with a Phred score below Q20. The data generated from this study was submitted to NCBI BioProject ID: PRJNA1178603. Cleaned reads were mapped to the reference genome using TopHat v2.1.1, and Cufflinks v2.2.1 was used for transcript assembly and expression quantification. Genes with a Log2FC>1.0 were considered upregulated, and those with Log2FC<1.0 were downregulated. DEGs were identified based on statistical significance (P < 0.05) and a false discovery rate (FDR < 0.05). The gene ontology (GO) enrichment of DEGs (Log2FC>1.0) was performed using the AgriGO v2 analysis tool kit using Bonferroni multiple testing with FDR < 0.05 (Tian et al., 2017) and the GO enrichment plots were generated using ggplot2 in R. Plant TFs were predicted from DEGs using iTAK program version 1.7 (https://itak.feilab.net/) with default parameters (Zheng et al., 2016). For the Heatmap generation, the normalized Log2FC was plotted using TB Tools software (Chen et al., 2020).

### Arabidopsis complementation and experimental conditions

The *TaMCO3* ORF (2019 bp; *TraesCS5A02G286900*) was cloned into the plant expression vector pGWB520 by using gateway cloning method. The resulting construct was introduced into Arabidopsis *atmco3* mutant lines (SALK_039183) through *Agrobacterium tumefaciens*-mediated transformation using the GV3101 strain. Following transformation, T_1_ generation seeds were screened for hygromycin resistance (20 μg/ml), and true transformants capable of reaching the four-leaf stage were selected. These plants were then transferred to soil pots and allowed to grow to maturity. T_2_ generation seeds were harvested, and the subsequent T_2_ plants were cultivated to obtain T_3_ seeds. Upon confirmation of successful transformation, Arabidopsis seeds from Col-0, *atmco3*, and TaMCO3-complemented lines were germinated and grown in Hoagland media for 5 days under control conditions. For phenotypic assays the seedlings were transferred to three different Fe treatments for 8 days: (i) with 80 µM FeEDTA (+Fe), (ii) without Fe and with 300 µM ferrozine (–Fe), and (iii) with 300 µM FeEDTA (+++Fe) to induce Fe toxicity. All the assessment and processing of the samples were done after 7 days of growth on the indicated plates.

### Perls and ferrozine staining in roots

Wheat (*Triticum aestivum L.*) or Arabidopsis seedlings were grown in a controlled environment using Hoagland’s nutrient solution, and supplemented with 40 µM Fe_2_SO□ and 60 µM Na-EDTA to maintain optimal Fe supply. Root samples were collected to assess Fe²□ accumulation using ferrozine staining and Perl’s Prussian blue staining. For Fe²□ detection, the roots were incubated in a freshly prepared 100 µM ferrozine solution. To ensure effective penetration, the samples underwent vacuum infiltration for 1 hour in complete darkness, minimizing the potential degradation of the ferrozine-Fe²□ complex. Following infiltration, roots were thoroughly rinsed with deionized water to remove excess stain and examined under a stereomicroscope for Fe²□ localization. In parallel, Perl’s Prussian Blue staining was employed to detect Fe³□ ions. Root samples (from wheat or Arabidopsis) were immersed in a solution containing 5% potassium ferrocyanide and 5% HCl and vacuum infiltrated under dark conditions for 1 hour to enhance reagent penetration. After staining, roots were washed with deionized water to remove any unbound stain, and the presence of Fe³□ was visualized by the formation of a blue precipitate. Following this, for DAB staining of wheat roots a thorough washing with distilled water was done to remove any excess staining (Perl’s) solution. Furthermore, the plants were kept in freshly prepared solution composed of 0.1% DAB (3,3’-Diaminobenzidine) in 0.1 M Sodium citrate buffer, pH 3.8 under dark conditions. A final rinse with distilled water was done to remove any residual dye. All DAB, ferrozine- and Perl-stained roots were examined and imaged under a stereomicroscope to document the distribution of Fe²□ and Fe³□ ions across different root zones.

### Yeast complementation assay

A yeast complementation assay was performed using the *S. cerevisiae* mutant strain *fet3fet4*, deficient in Fe uptake (Spizzo et al., 1997). The coding sequences of multiple wheat TaMCO3 were cloned into the pYES2 expression vector (at site *BamH*I and *EcoR*I present at MCS), generating two constructs: pYES2-TaMCO3 and pYES2-PMTaMCO3 (the latter containing the plasma membrane-PM domain of yeast *FET3* fused with GFP). The FET3 PM domain was inserted using overlapping primers in Table S8. These constructs were transformed into *S. cerevisiae* wild-type BY4741 and *fet3fet4* strains via the lithium acetate method. Transformed cells were selected on SD -Ura medium and their growth was assessed by spotting serial dilutions onto DOB -Ura plates under two conditions: +Fe (50 mM MES and 100 µM ferric citrate) and –Fe (50 mM MES only). The assay evaluated the ability of *TaMCO3* to complement Fe uptake deficiency in *fet3fet4* mutants. For GFP detection yeast strains, images were captured in replicates at 25X using an Zeiss LSM 880 Confocal Laser Scanning Microscope.

### In-silico identification of wheat MCOs genes and phylogenetic analysis

To analyse the phylogenetic relationship between yeast *fet3* and wheat, *Arabidopsis, O. sativa* MCOs, and proteins sequences were used. These MCOs protein sequences were extracted from Ensembl Plants by doing fet3 protein sequence BLAST with wheat, *Arabidopsis* and *O. sativa* whole genome. In total 77 genes from *Triticum aestivum*, 12 from *Arabidopsis thaliana* and 12 from *Oryza sativa* were extracted from Ensembl Plants. The putative IDs and protein sequences were then aligned using the MUSCLE algorithm (Edgar, 2004) and a rooted tree was constructed using the maximum likelihood method using MEGAX software (Kumar et al., 2018). The phylogenetic tree was constructed to depict wheat, *Arabidopsis* and *O. sativa* MCOs which have the most similarity with yeast *fet3.* Twenty-nine high-confidence TaMCO proteins were divided into six clades on the basis of their sequence similarity. Gene structure for the TaMCO transcripts were then analysed to determine their conserved motif architecture. These conserved motifs were predicted using MEME (Bailey et al., 2015). A total of 8 conserved motifs designated as motifs 1-8 were identified with varying lengths. Motifs 1,2,3,6 were conserved among yeast *fet3* and TaMCOs. These motifs will be further subjected to more *in-silico* validation. Along with this, the arrangement of introns and exons of wheat MCOs and their homeologs was analysed by using the Gene Structure Display Server (GSDS) (Hu et al., 2014).

### RNA extraction and gene expression analyses

Total RNA extractions were carried out using Trizol Reagent (Rio et al., 2010). RNA quantity and purity were assessed using a NanoDrop™ Lite Spectrophotometer (ThermoFisher Scientific). To eliminate genomic DNA contamination, RNA samples underwent DNase treatment using TURBO™ DNase (Ambion, Life Technologies). For qRT-PCR analysis, cDNA was synthesized from DNase-treated total RNA using the Invitrogen SuperScript III First-Strand Synthesis System SuperMix (ThermoFisher Scientific). Gene expression quantification was performed with the QuantiTect SYBR Green RT-PCR Kit (Qiagen, Germany) on a Bio-Rad CFX96 Real-Time PCR detection system. Gene-specific primers, designed to amplify a 150-250 bp region from all three homologs of the wheat genes, were generated using Oligocalc software. Primer sequences used in qRT-PCR are listed in **Table S8**. Relative transcript levels were calculated using the 2−ΔΔCT method (Livak and Schmittgen, 2001), with the ADP-ribosylation factor gene (TaARF) serving as an internal control.

### Sub-cellular localization of TaMCO3

*TaMCO3* amplification was performed from pJET1.2 using gene-specific primers and subsequently ligated into the pENTR™/D-TOPO™ vector. Following sequence confirmation, an LR reaction was conducted with the positive TaMCO-pENTR TOPO and the pSITE-cEYFP vector. The resulting TaMCO3-cEYFP construct, along with the pSITE-cEYFP empty vector and the plasma membrane marker clone pMRK-RFP, was transformed into Agrobacterium strain GV3101. The transformed GV3101 cells were resuspended in an agroinfiltration medium [10 mM MES (pH 5.6), 10 mM MgCl2, and 200 μM acetosyringone] and used to inoculate the abaxial surfaces of six- to eight-week-old *Nicotiana benthamiana* leaves at an OD600 of 0.6-0.8 using syringe infiltration. The leaf tissues were visualized for fluorescent signals 48 hours post-infiltration using a LEICA DM600CS confocal microscope. YFP fluorescence was detected at 500–530 nm, and red fluorescence (RFP) was detected at 580–630 nm.

### Wheat transformation, screening and immunolocalization of TaMCO3

To generate *TaMCO*3 overexpression lines in the wheat variety ‘Fielder,’ Agrobacterium-mediated transformation was conducted as previously described (Hayta et al., 2021). The pGWB520 vector containing *TaMCO3* driven by the constitutive CaMV35S promoter was introduced into *Agrobacterium* strain AGL-1, and positive colonies were selected. Immature wheat embryos were transformed with the construct. Transformed calli were selected on hygromycin at 50 mg/L for the first round of selection, 30 mg/L for the second, and 15 mg/L during the regeneration phase. The presence of the transgene in the regenerated plants was confirmed using PCR analysis (Hygromycin/M13 Primers), following the protocol of (Bhati et al., 2016).

For whole-mount immunolocalization and confocal microscopy wheat root samples were fixed in methanol for 20 min. at 37°C. Immunolocalizations of sectioned wheat root were performed essentially as described earlier (Pasternak et al., 2015). The primary antibody (anti-c-Myc tag Mouse monoclonal antibody Catalog no. M1001050; Thermofisher, USA) was diluted 1:100 in Blocking buffer (containing 2% bovine serum albumin in 1xMTSB buffer), and the secondary antibody (Alexa Flour^TM^ Plus 488 anti-mouse Catalog no. A32723; Thermofisher, USA) was diluted 1:800 in blocking buffer. Antibody-treated sections were transferred to microscopic slides and observed with the confocal microscope. Images were captured in replicates at 10X using an Zeiss LSM 880 Confocal Laser Scanning Microscope. To measure the degree of chlorosis, the fully expanded youngest leaves of seedlings were used, with 3-4 spots per leaf. The measurements were taken using a portable SPAD metre (Konica Minolta SPAD-502, Japan), and n=36 seedlings were used for each treatment.

### Ferroxidase assay in wheat roots

For ferroxidase detection *Triticum aestivum* cv. C306 seeds were grown in control and Fe-deficient media for 8 days and 14 days. Total protein was isolated from roots by using HEPES buffer and PMSF, and protein concentration was then quantified by using Bradford reagent protein estimation. Equal amounts of protein were used to perform the ferroxidase assay. The oxidation of Fe (II) was assayed by monitoring the rate of disappearance of ferrous ammonium sulfate using the ferrous chelator ferrozine. Reactions were carried out in microcentrifuge tubes containing 1050 µl buffer (450 mM Na-acetate (pH 5.8), 100 µl CuSO_4_) and 15 µl of total protein from roots (2 µg/ml). After which the reaction was initiated by adding 225 µl substrate (Fe(II)SO_4_, 357 µM) containing 100 µM CuSO_4_ (Müller et al., 2015). The aliquots (200 µl) were then removed at appropriate intervals and transferred to 96 well microtiter plates for reaction-quenching with 14µl of 18mM ferrozine. The rate of Fe^2+^ oxidation for the Fe^2+^-ferrozine complex was calculated from the decreased absorbance at 560nm (Hoopes and Dean, 2004).

## Acknowledgements

The authors thank the Executive Director, NABI for the facilities and support. This work was supported by the NABI-CORE grant to AKP. DBT-eLibrary Consortium (DeLCON) is acknowledged for providing timely support and access to e-resources for this work.

## Contributions

Conceptualization: AKP, TL, SS and RJ; Project administration: AKP, JK; Investigation: RJ, DT, GS, M, SS, VS; Formal analyses: RJ, VS, ER, VM, DT, GS, HB; Visualization: RJ, KA, VM, TL, Resources: SS, TL Methodology: All authors. Software: GS, VM, RJ; Validation: RJ, GS, DT, KA. Supervision: AKP, TL, SS, ER. Writing-Original draft: RJ, AKP, GS, KA; Writing-Review and Editing: TL, ER, SS, RJ; Funding Acquisition: AKP, TL.

## Data Availability

The data generated from this study have been deposited in the NCBI Sequence Read Archive (SRA) database and are accessible with the submission ID SUB14812731 with BioProject: PRJNA1178603.

## Supplementary Tables

Table S1: –Fe induced DEGs analysed in the wheat root tip

Table S2: GO enrichment analysis of 8 day -Fe DEGs detected within the wheat root tip

Table S3: GO enrichment analysis of DEGs common between 8-day whole root and Root Tip under –Fe

Table S4: GO enrichment of A, B, and D genome 8 day root tip DEGs under Fe-deficient conditions.

Table S5: GO enrichment of wheat MCOs DEGs from root tip under –Fe.

Table S6: List and GO enrichment of Root Tip Specific Marker Genes.

Table S7: List and GO enrichment of Cluster 4 Pericycle DEGs. Table S8: List of the primers used in this study.

