## Supplementary material for "Integrative spatial transcriptomic analysis pinpoints the role of the ferroxidase, TaMCO3, in wheat root tip iron mobilization": data

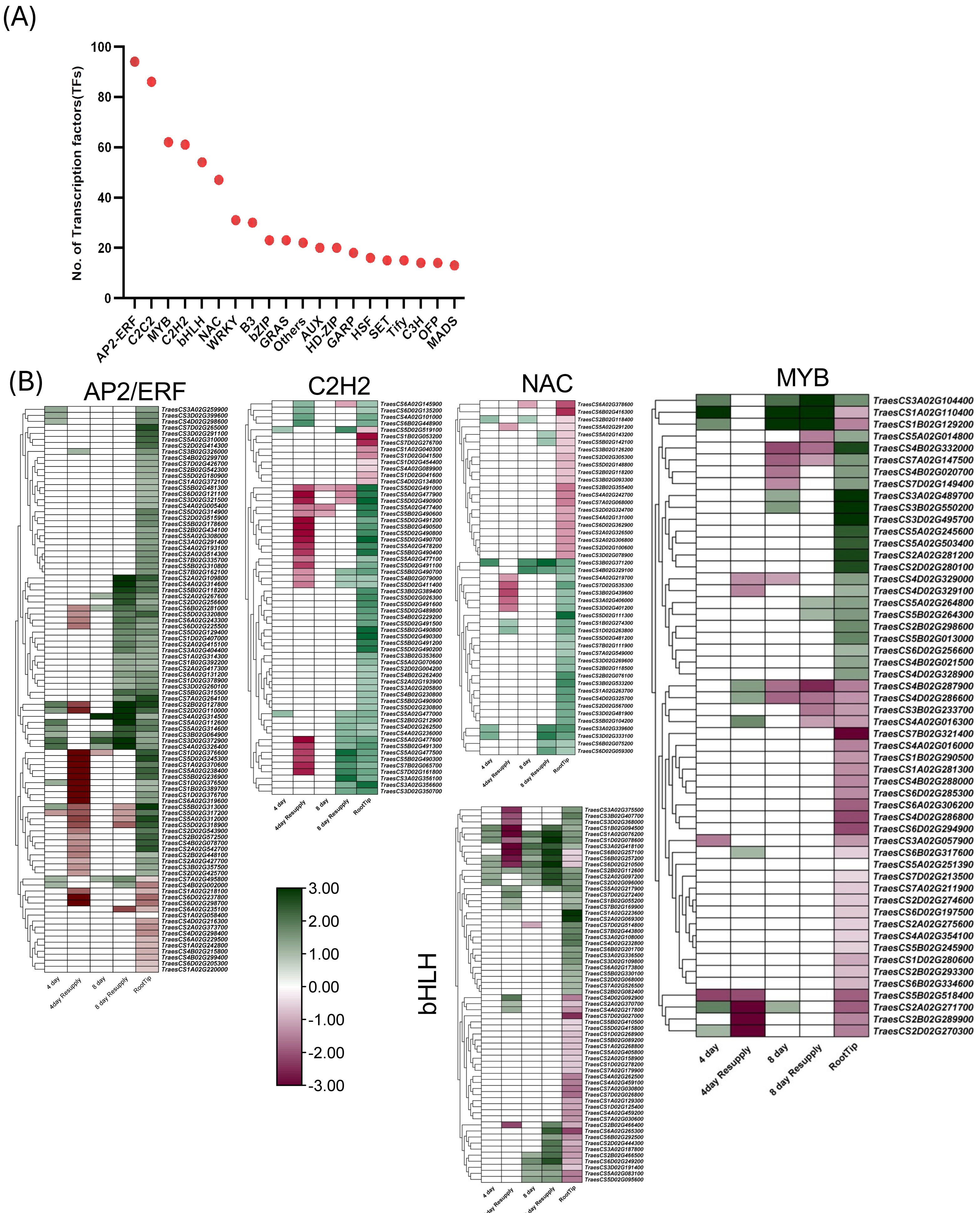


Figure S1: (A) number of transcription factors (TFs) accounting to different family of genes. (B) Heatmap analysis of different TFs during –Fe.

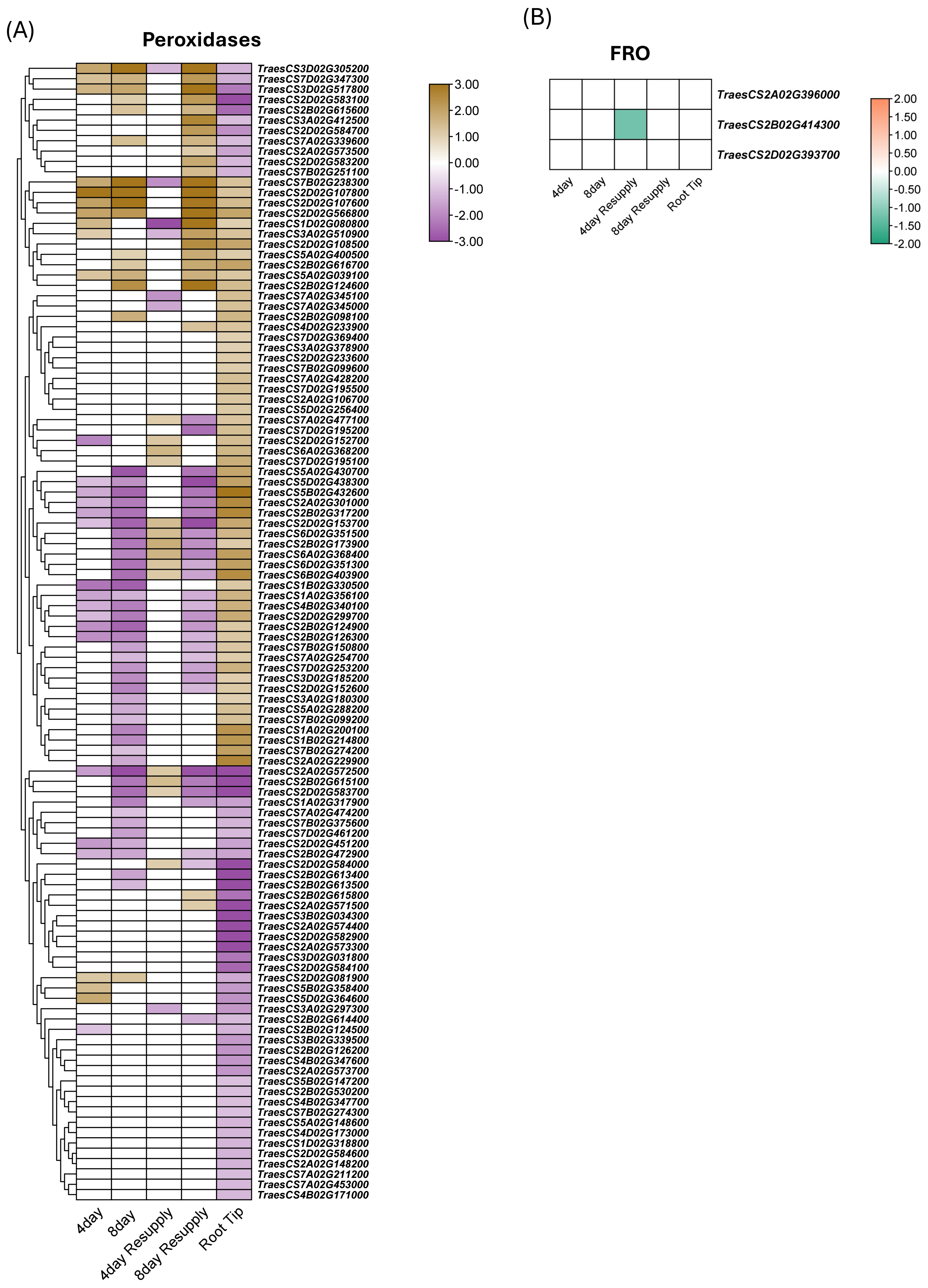


Figure S2: Heatmap analysis of (A) peroxidases family genes and (B) FRO2 (ferric chelate reductase) genes under -Fe

| Percent Identity Matrix | TraesCS4D02G035000 | TraesCS7B02G202500 | ScFET3 | TraesCS5A02G286900 | TraesCS7D02G176300 | TraesCS1D02G044300 | TraesCS6D02G361500 | TraesCS3A02G452600 |
|---|---|---|---|---|---|---|---|---|
| **TraesCS4D02G035000** | 100 | 19.48 | 18.01 | 21.09 | 22.25 | 18.33 | 21.98 | 21.48 |
| **TraesCS7B02G202500** | 19.48 | 100 | 19.81 | 28.99 | 23.34 | 24.9 | 25.93 | 26.44 |
| **ScFET3** | 19.81 | 19.81 | 100 | 29.07 | 26.19 | 26.18 | 25.96 | 22.54 |
| **TraesCS5A02G286900** | 21.09 | 28.99 | 29.07 | 100 | 33.15 | 31.19 | 32.35 | 30.98 |
| **TraesCS7D02G176300** | 22.25 | 23.34 | 26.19 | 33.15 | 100 | 43.11 | 40.63 | 37.19 |
| **TraesCS1D02G044300** | 18.33 | 24.9 | 26.18 | 31.19 | 43.11 | 100 | 47.55 | 47.87 |
| **TraesCS6D02G361500** | 21.98 | 25.93 | 25.96 | 32.35 | 40.63 | 47.55 | 100 | 57.8 |
| **TraesCS3A02G452600** | 21.48 | 26.44 | 22.54 | 30.98 | 37.19 | 47.87 | 57.8 | 100 |

Figure S3:Percentage identity matrix comparing *Saccharomyces cerevisiae* ferroxidase *ScFET3* with wheat multicopper oxidases (*TaMCOs*) expressed under iron-deficient (-Fe) conditions. The matrix highlights the sequence identity among these proteins, reflecting potential functional similarities

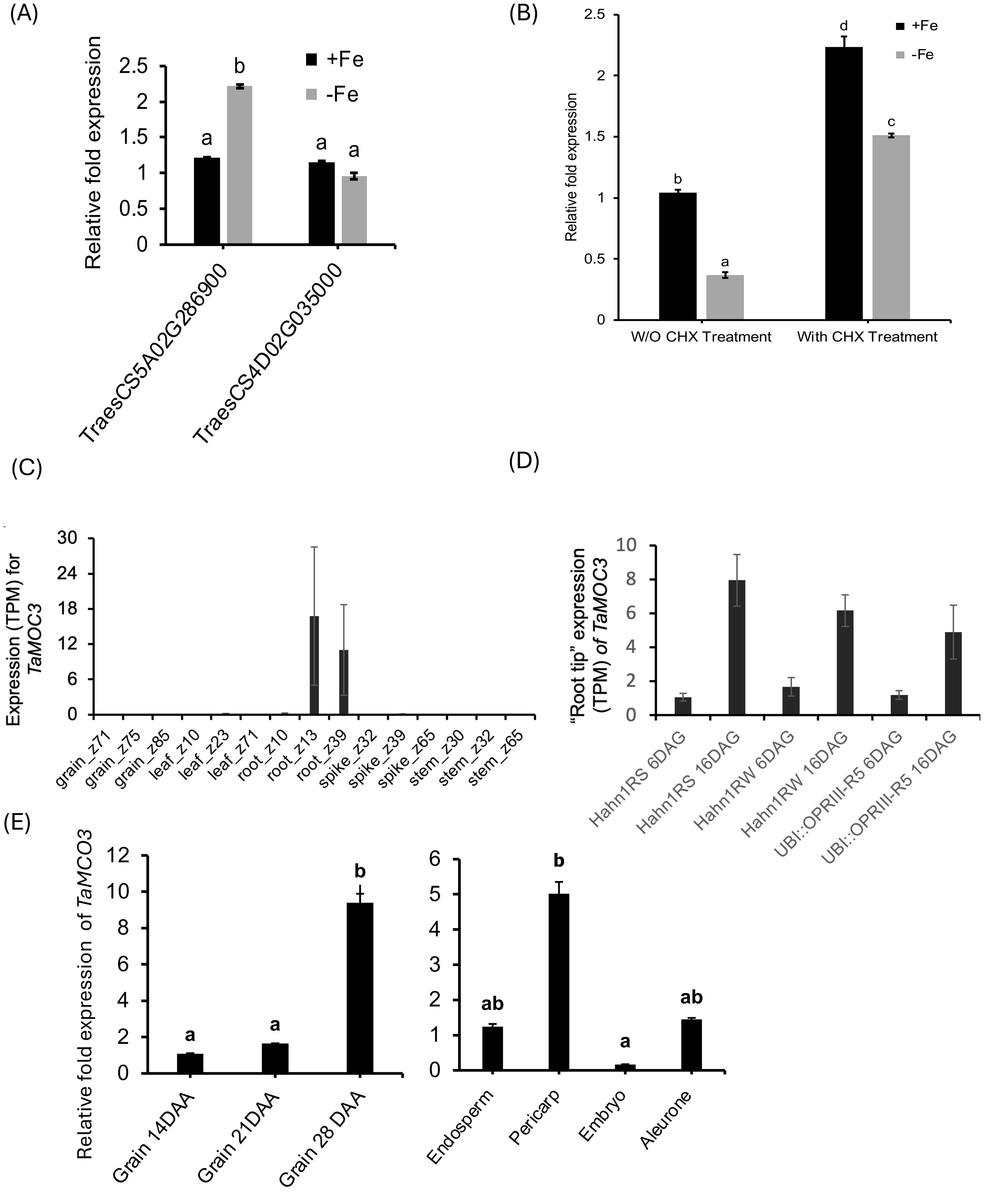


Figure S4: (A) qRT-PCR analysis of selected wheat *MCO* in root tips under iron-sufficient (+Fe) and iron-deficient (−Fe) conditions. Gene expression levels were normalized to *TaARF1*, and relative fold changes were determined using the $2^{-\Delta\Delta CT}$ method. Error bars represent standard deviation (n = 9), and different letters above the bars denote statistically significant differences ($P < 0.05$) as determined by two-way ANOVA..(B) qRT-PCR expression analysis of *TaMCO3* under +Fe and −Fe conditions with or without cycloheximide (CHX) treatment (50 µM, 1 h) to inhibit protein translation (n=6). (C) **(C)** *In-silico* expression analysis of *TaMCO3* in different tissues during the developmental time course of plant development. **(D)** Gene expression analysis of *TaMCO3* in different genetic backgrounds as reported by (Gabay et al., 2023). **(E)** qRT-PCR analysis of *TaMCO3* during grain development (days after anthesis-DAA) and in different grain tissue as mentioned.

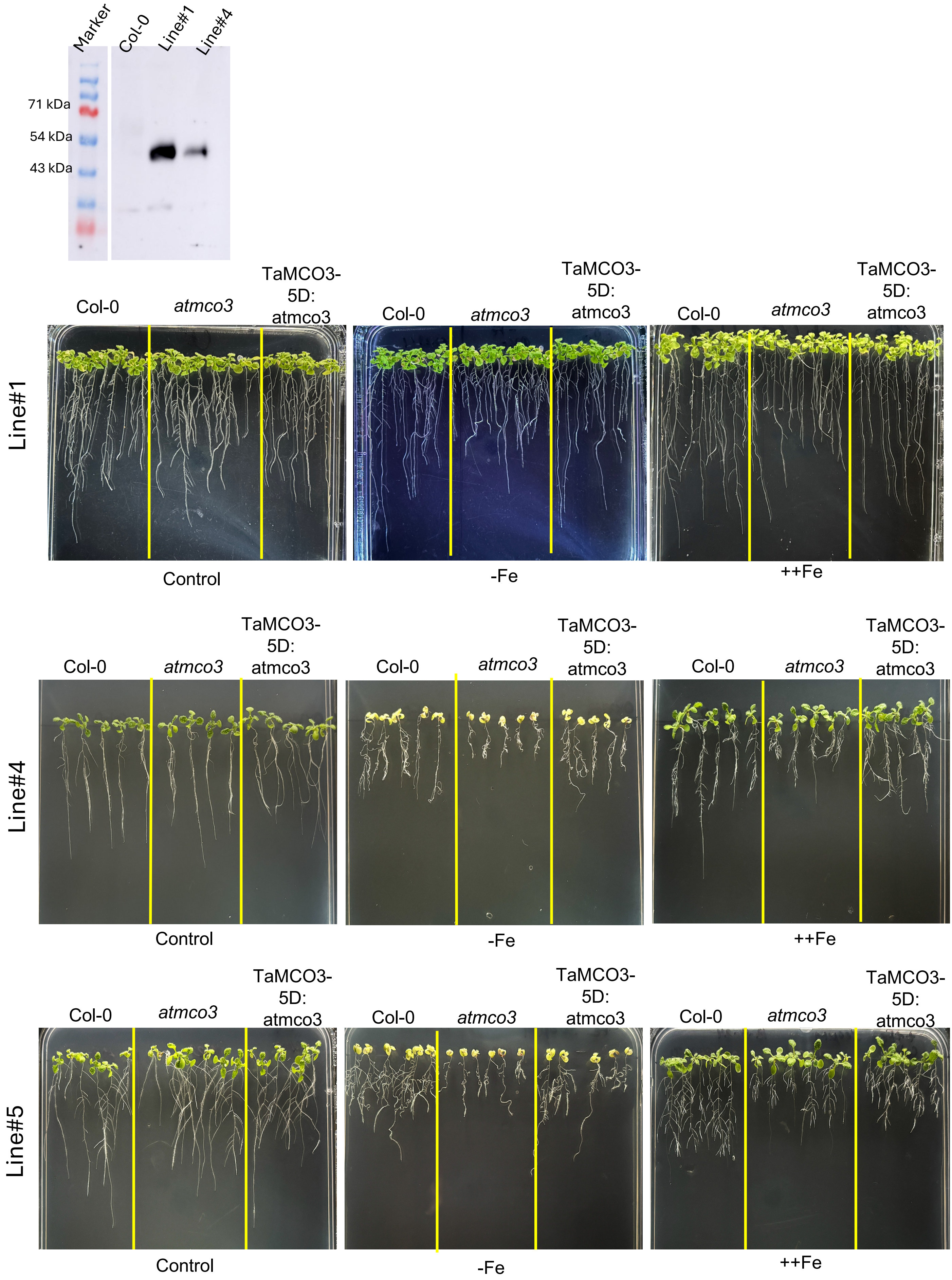


Figure S5: Western analysis of complemented lines in Arabidopsis and its characterization. Characterization of atmco3 complementation with TaMCO3 under –Fe (1 μM), ++Fe (excess 300 μM) and control condition (80 μM).

(A)

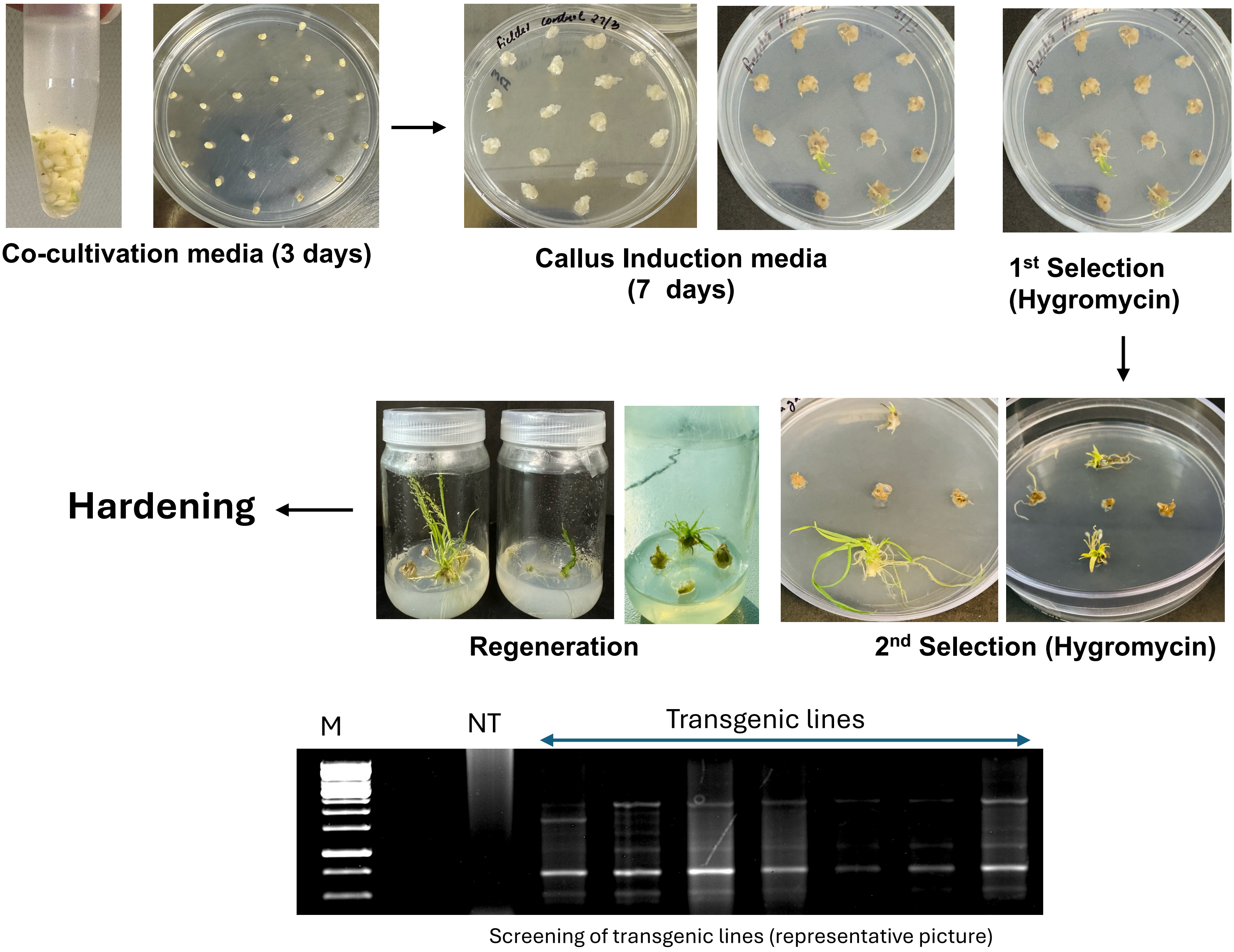


(B)

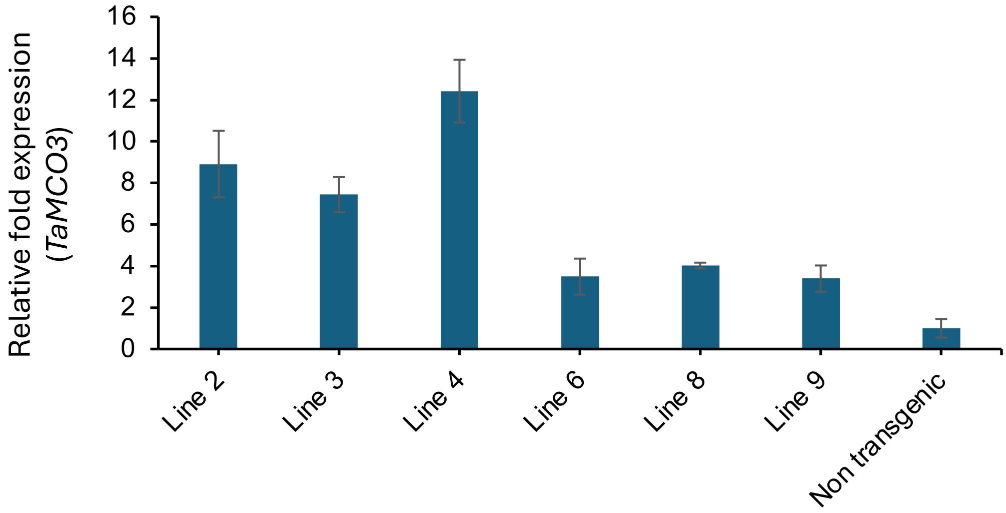


Figure S6: (A) Representative pictures for differ stages of Wheat transformation and screening process to generate TaMCO3 overexpressing lines in cv. Fielder. (B) Relative fold expression of TaMCO3 in multiple wheat transgenic lines.

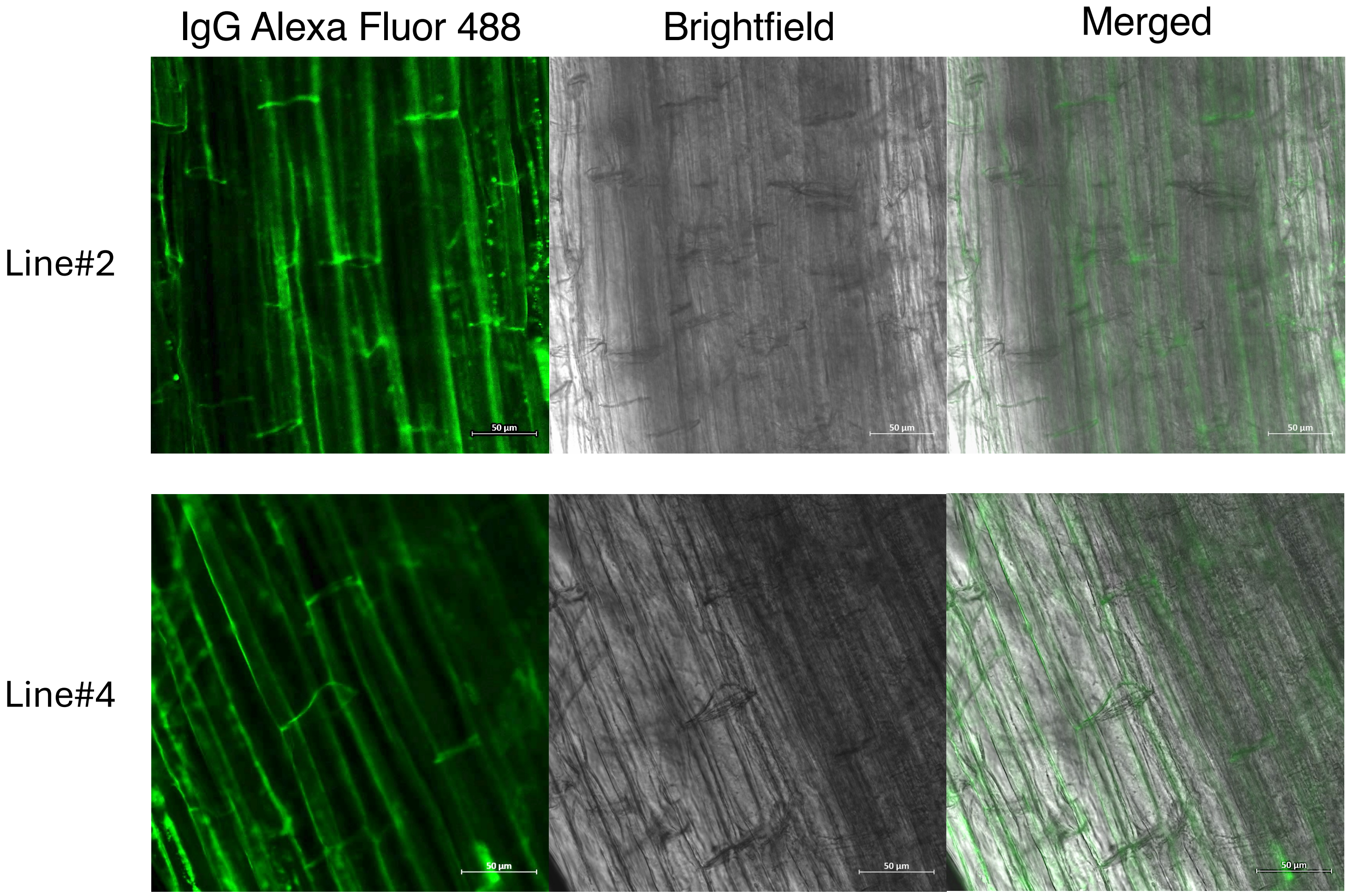


Figure S7: TaMCO3 overexpression line whole-mount immunolocalization of leaf tissue, antibody-treated sections were transferred to microscopic slides and observed with confocal microscope. Images were captured in replicates at 10X using an Zeiss LSM 880 Confocal Laser Scanning Microscope.

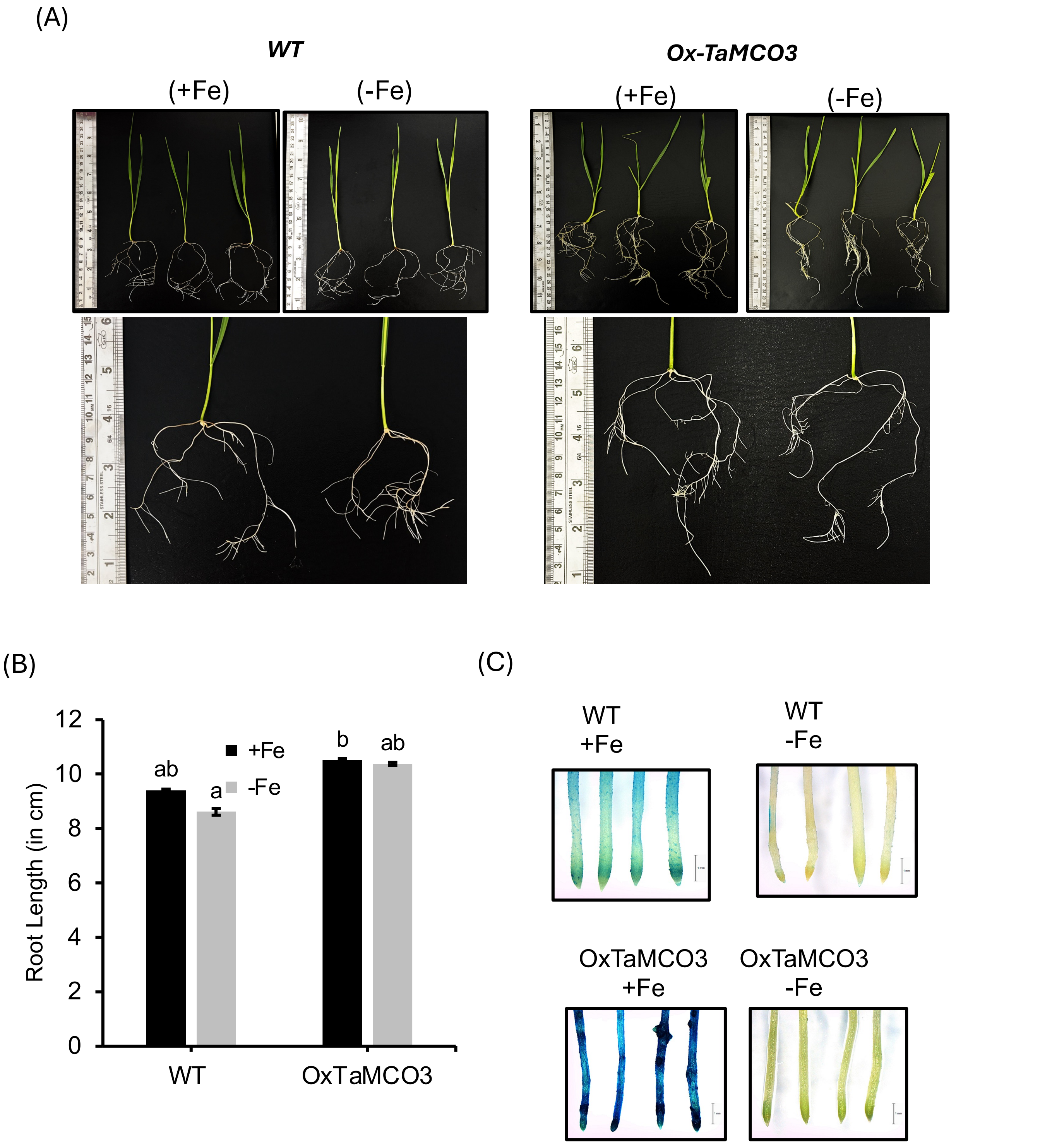


Figure S8: (A) Phenotypic response of wheat overexpressing TaMCO3 and non-transgenic lines under –Fe and control conditions. (B) Measurement of root length (n=3) of wheat seedlings subjected to +Fe –Fe. (C) Perls stain of the roots from the above experiments to check the accumulation of Fe.

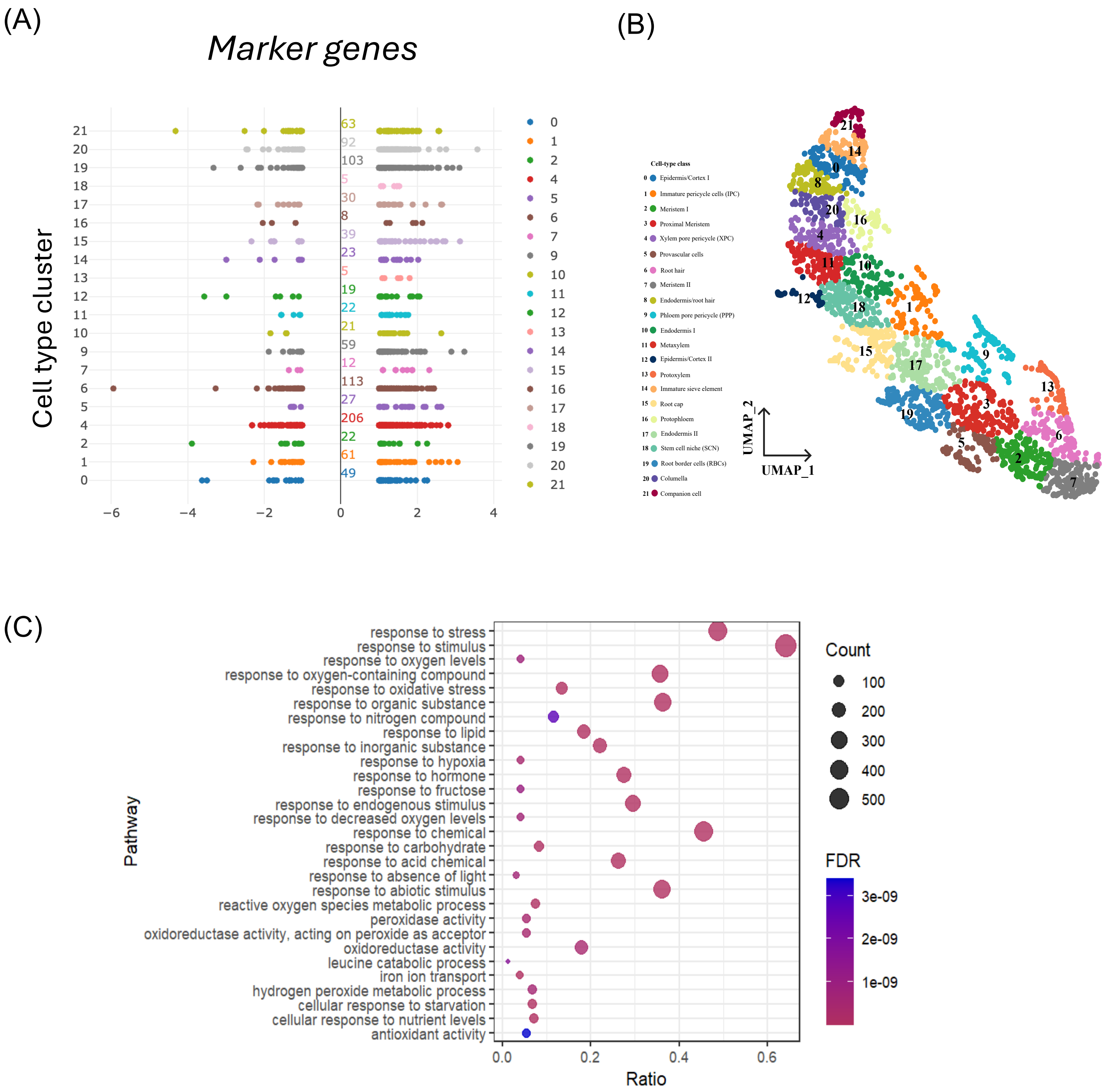


Figure S9: Categorization of differentially expressed genes (DEGs) associated with iron deficiency based on their expression patterns across distinct cell types, as analysed by snRNA-sequencing of wheat root tips (Zhang et al., 2023) . (A) A scatter plot shows the correlation of DEGs expressed under Fe deficiency with their expression levels across various cell types. (B) UMAP visualization demonstrating the spatial distribution of –Fe DEGs across different cell-type clusters as identified in wheat root tips. Each data point represents the specific DEG expressed within a single cell type, with colours denoting the respective cell types. (C) Analysis of enriched Gene ontology terms for root specific marker genes by creating dot bubble plot of enriched GO categories.

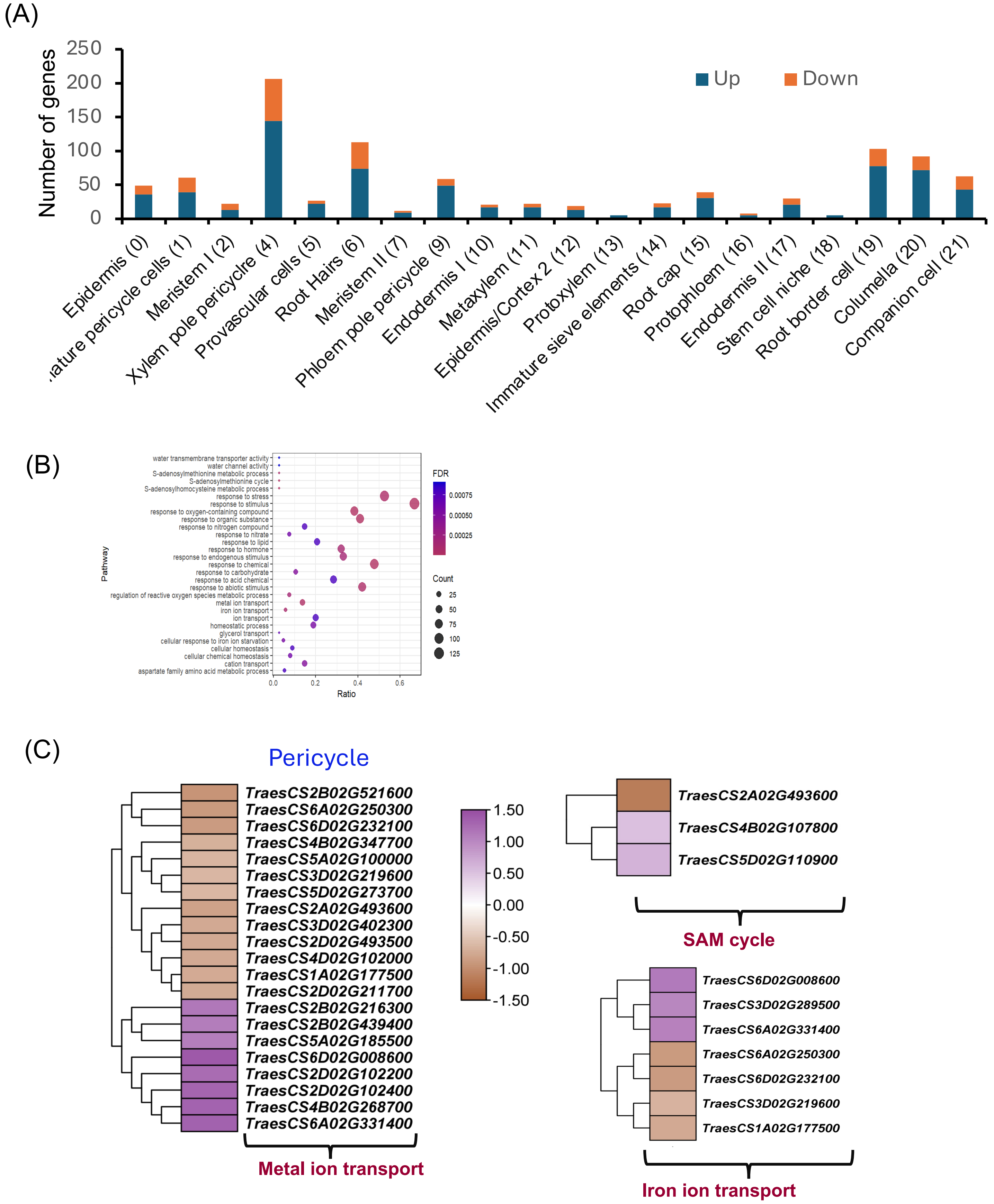


Figure S10: Cell-type distribution of the DEGs and their GO annotations. (A) The graph shows the number of genes that were differentially upregulated in root tip –Fe RNA-seq data across various cell type regions. The data highlight the gene expression changes observed in response to iron deficiency within specific root single cell types. (B) GO enrichment analysis was performed for various overlayed DEGs on the single-cell type cluster of pericycle, revealing different pathways associated with upregulated genes. The results illustrate the distinct pathways enriched in this cell type in response to the root tip under Fe deficient conditions. (C) Heat map analysis to visualize differentially expressed genes (DEGs) that were regulated in pericycle highlighting the differential regulation of genes in response to –Fe.
